# Population geometry reveals directed coupling and transient bistability in spontaneous pituitary secretion

**DOI:** 10.64898/2026.04.05.716480

**Authors:** Ana Aquiles, Aparicio Arias Juri, Lafont Chrystel, David J. Hodson, Santiago-Andres Yorgui, Patrice Mollard, Tatiana Fiordelisio

## Abstract

The pituitary gland operates as an organized signaling network in which endocrine cell populations coordinate hormone secretion, through homotypic and heterotypic interactions, yet the contribution of spontaneous intrinsic activity in shaping population-level dynamics remains poorly understood. Using geometric analysis of population trajectories — including subspace alignment, manifold separation, and directed coupling metrics — we identified two classes of spontaneous oscillatory signals associated with distinct cell populations exhibiting asymmetric geometric dominance and a reproducible temporal lag. Our results support that spontaneous activity generates a self-sustained oscillator exhibiting transient bistability, linked to increased physiological demand, with slow oscillations reflecting the properties of an excitatory resonator capable of self-oscillating dynamics without external drive. A low-rank recurrent neural network model recapitulated the empirical geometric landscape under three coupling conditions, confirming that directed population coupling underlies the observed coordination. These findings suggest that intrinsic population dynamics play a central role in coordinating pituitary secretion, with implications for understanding hormonal dysregulation in secretory adenomas and other pituitary disorders.

**Graphical Abstract:** 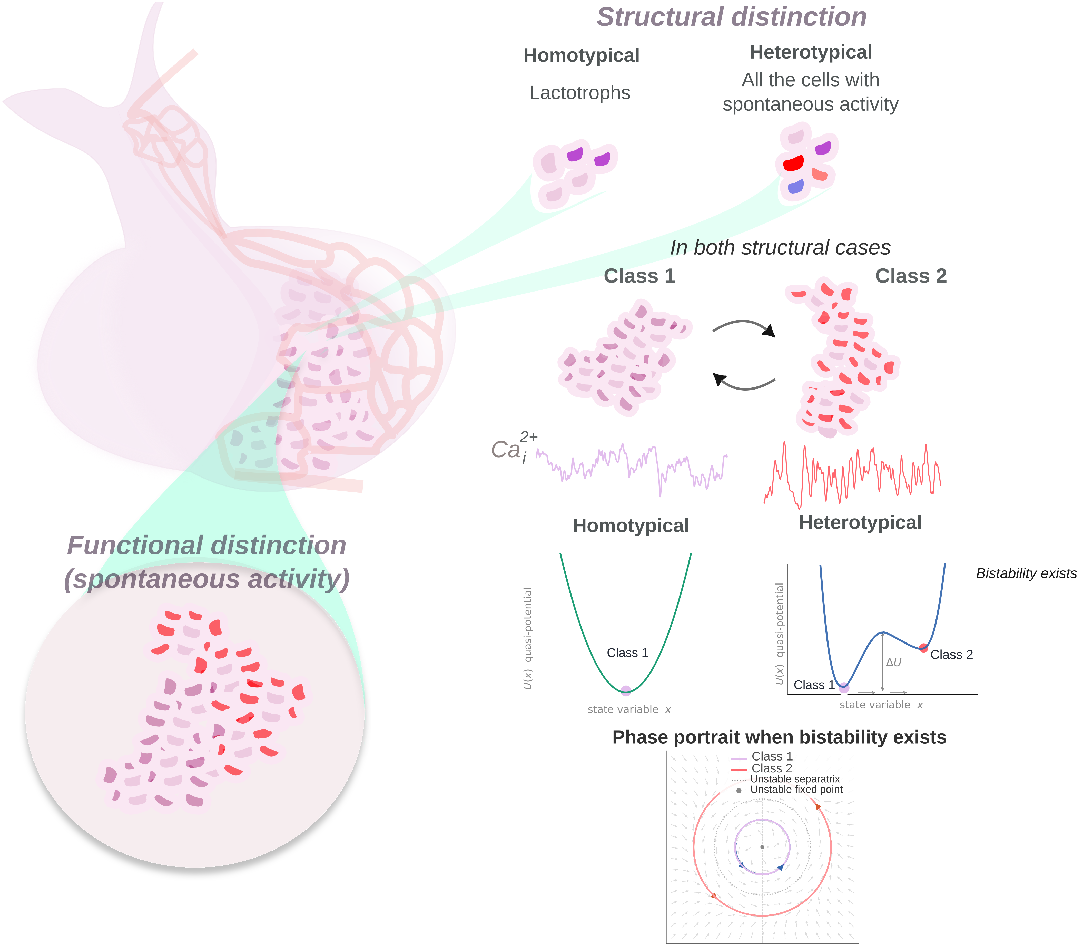

Structural and functional distinctions between homotypic and heterotypic interactions have been widely described in the pituitary endocrine system. However, whether functional differences in intrinsic calcium time-series dynamics are relevant to pulsatile hormone secretion remains unexplored. Here, we classify the spontaneous activity underlying both homotypic and heterotypic interactions and characterise their synchrony. We find that heterotypic interactions exhibit transient bistability, consistent with a Hopf-type oscillator regime, in which slow oscillations drive secretory output according to physiological demand.

## 1 Main

Pituitary cell communication is widely recognized as a network that is organized and an integral sum of the interactions involved in sending and receiving signals among its diverse cellular composition [1]. This communication improves functional outcomes and highlights its crucial role in efficient functioning and adaptation [2]. Even so, our understanding of how cell signaling patterns contribute to spontaneous endogenous secretion and plasticity remains limited. It is essential to investigate spontaneous activity and its influence on homotypic interactions (within the same endocrine cell group) versus heterotypic interactions (between different endocrine cell groups). These differences in signaling could be the primary responsible for the variability in activation oscillations when responding to physiological challenges [3-5]. For example, in pathological conditions, non-functioning pituitary adenomas are often referred to as “silent” because they do not produce significant levels of hormone [6]. However, there are also adenomas that result in hormonal overproduction [7]. The origins and causes of these functional differences are not yet fully understood. Therefore, gaining insight into the role of spontaneous signal transmission in triggering secretion may help clarify why some conditions remain non-secretory, while others lead to hormonal secretion in both health and disease.

Research consistently shows that pituitary endocrine cells communicate through multiple signaling modes [2, 5, 8-12], all of which are modulated by external stimuli — such as hypothalamic hormones or downstream hormonal regulation. Although spontaneous activity has received comparatively less attention, two endocrine cell populations — lactotrophs and somatotrophs — exhibit substantial spontaneous activity, allowing them to secrete hormones independently of external triggers [13-15]. Understanding how these cell groups integrate information differently is essential, as this process is shaped by the architecture of their communication networks [5, 8]. Within the homotypic lactotroph network, this structure underlies key physiological adaptations, including the reorganization of lactotroph signaling connectivity during lactation and pregnancy [5] as well as structural changes in cell density and spatial arrangement [5, 16]. Furthermore, evidence supports the existence of heterotypic communication and coordination among distinct endocrine cell groups, equipping the gland to generate unified responses to evolving physiological demands demands[17]. Yet no direct comparison exists between the population dynamics governing homotypic coordination and those potentially underlying heterotypic integration. This is particularly relevant in light of reported homotypic coordination during lactation and female reproductive plasticity [5], which suggests that broader heterotypic interactions may also contribute to shaping these or related physiological states. Beyond homotypic interactions, signal transmission may engage multiple endocrine cell types, shaping the systemic mechanisms through which cells spontaneously exchange signals.[3, 4, 13]. In this study, we identified two classes of spontaneous signals with distinct oscillatory dynamics — slow and fast — reflecting different modes of activity within the gland. Leveraging the well-characterized spontaneous secretion of lactotrophs [5], we then asked how their population-level coordination changes under conditions that recapitulate heterotypic interactions, assessing both network coordination and the geometrical structure of population dynamics across homotypic and heterotypic configurations. Our findings revealed that during spontaneous activity, a self-sustained oscillator is generated, resulting in momentary bistability in cell organization that is associated with an increase in physiological demand. This mechanism may explain how population leaders coordinate within the geometric structure of the cell networks. We show that slow oscillations predominate across both homotypic and heterotypic interactions, reflecting the computational properties of an excitatory resonator that sustains self-oscillating dynamics. Such a system retains the flexibility to accommodate heterogeneous inputs while generating signaling patterns aligned with physiological demands. Our findings further underscore the dual role of excitatory and inhibitory-like activities in regulating spontaneous coordination during heterotypic integration — dynamics that may represent the scaffold underlying the generation of pulsatile secretion. Together, these results deepen our understanding of spontaneous pituitary signaling and carry potential implications for conditions such as secretory tumors and other hormonal disorders.

## 2 Results

### 2.1 Spontaneous calcium signals wave features in homotypical and heterotypical interactions

To characterize the temporal wave signatures observed during spontaneous activity in both homotypic and heterotypic interactions, we reanalyzed previously acquired calcium imaging recordings from lactotrophs obtained under four main physiological conditions, basal hormonal stimulation, (nullipara) basal condition,(primipara) lactation, weaning, and multipara states. Additionally, we expanded this analysis to evaluate the overall population activity across these conditions. These physiological states are associated with sustained long-term integration and cell-network coordination in response to increased physiological demands (Figure 1) [5, 18, 19]. Spontaneous calcium traces display variability in their waveforms, regardless of their physiological effects. However, the type of interaction influences these wave shapes, as illustrated in Figure 2A. Both types of interactions exhibit a signal coupling pattern characterized by a low-frequency component paired with a higher-frequency component. This observation motivates us to quantify the Phase-Amplitude Coupling between these two frequency ranges by calculating the modulation index, which measures the strength of the coupling. As shown in the representative example (Figure 2A, top right panel), the lowest phase frequencies exhibit modulation across a wide range of amplitude frequencies in both homotypic and heterotypic spontaneous activity.

**Figure 1:**
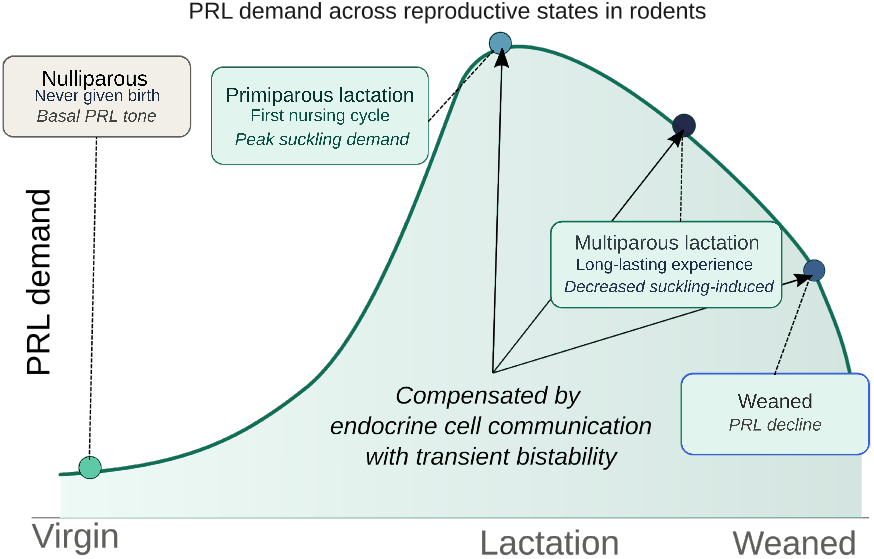
Hypothesized prolactin (PRL) hormonal demand across reproductive states in rodents, under which the physiological interpretation of our result were based. Based on prior research on prolactin secretion dynamics in rodents, we hypothesize that PRL demand follows the trend depicted in the diagram: baseline levels correspond to Nullipara animals 53]; a peak occurs in primiparous dams (first lactation) driven by the suckling-induced neuroendocrine reflex 5, 54, 55]; PRL demand then partially declines in multiparous dams due to reproductive-experience-induced adaptation of central PRL feedback sensitivity 5, 56]; and demand declines further in the weaned condition 53, 57], when PRL is no longer required to sustain milk production.

**Figure 2:**
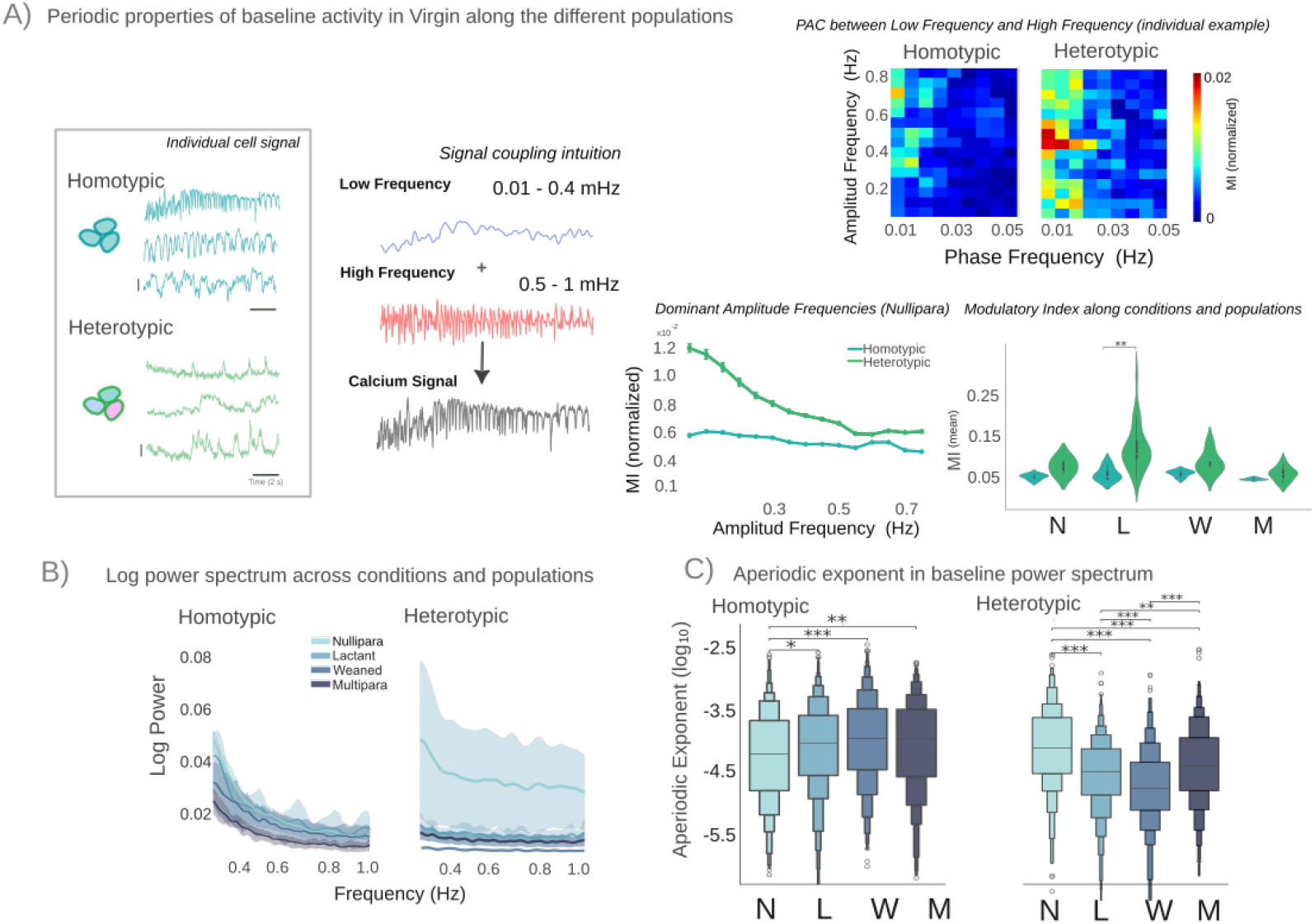
Periodic and aperiodic descriptors of spontaneous oscillatory activity across physiological conditions. **(A)** Spontaneous activity recorded from three distinct cell populations under control (Nullipara) conditions. Blue-green traces correspond to lactotrophs; green traces represent the full mixed population (see diagram, left panel). The right panel shows a representative phase−amplitude coupling (PAC) comodulogram in the Nullipara condition; the colour scale indicates the Modulation Index (MI) normalised within each cell population. Lower sub-panels show the dominant amplitude frequencies (left) and the MI compared across conditions and populations (right; group-level comparison using Kruskal-Wallis for Lactotrophs (*H* = 5.9, *p* = 0.12, n.s.) and pairwise Mann-Whitney U with Bonferroni correction for Heterotypic recordings; Heterotypic MI was significantly higher in Lactant than in Multipara (*p* = 0.003, ∗∗), with no other significant pairwise differences detected; within the Lactant condition, Heterotypic MI was significantly greater than Lactotroph MI (*p <* 0.05, ∗)) **(B)** Mean log-power spectrum for each population (dark line ± shaded variance), shown separately for homotypic (lactotroph) and heterotypic populations across all conditions. **(C)** Distribution of aperiodic exponent values across conditions and populations (Kruskal-Wallis + Dunn post-hoc with Bonferroni correction; ∗ = *p <* 0.05, ∗∗ = *p <* 0.01, ∗ ∗ ∗ = *p <* 0.001).

Group-level comparison revealed a wide range of dominant amplitude frequency modulation from 0.1 to 0.6 Hz (Figure 2A, right panel, green line), suggesting a broader adaptation in modulating different frequency values. In contrast, homotypic interactions show a notable increase at higher amplitude frequencies between 0.6 and 0.7 Hz, as indicated by the blue line. These differences reveal distinct modulation profiles for homotypic and heterotypic interactions, suggesting a preference for certain frequency values. Generally, homotypic interactions display lower modulation index values across various physiological states, with heightened variability observed during lactating and weaning conditions. A similar trend can be seen in heterotypic interactions, though they tend to have higher overall values.

To further analyze spectral organization, we quantified the features of the aperiodic exponent and measured the aperiodic exponent of the logarithmic frequency spectrum (see Figure 2B). The homotypic spectrum demonstrates a hierarchical order that aligns with the increasing hormonal demands of the physiological state [5]. In contrast, heterotypic interactions do not show this trend but exhibit greater variance in the spectrogram decay during nulliparous conditions. These findings are further supported by a direct estimation of the aperiodic exponent across the spectrum (Figure 2C).

Recent research on aperiodic spectral structure in other biological systems suggests that these features may reflect aspects of large-scale signal organization, including changes in rhythmic activity. In some contexts, such patterns have been discussed in relation to excitation-inhibition balance [20–22]. Lower aperiodic values, indicated by a flatter decay across frequencies, may suggest increased inhibitory activity within the system. Conversely, higher aperiodic values, characterized by a steeper frequency decay, may indicate a dominance of excitatory activity. However, it is important to note that these interpretations can vary depending on the specific system being analyzed. In our results, we will use these interpretations as a framework to investigate whether similar excitatory-inhibitory patterns can be identified in our findings. For instance, the variability observed in heterotypic interactions during nulliparous conditions indicates a broader range of spontaneous spectral states. In contrast, during lactation and weaning, both the variance and aperiodic values decrease, suggesting a more constrained spectral organization under these physiological conditions. This reduction may be related to significant hormonal shifts associated with the physiological state. For example, during lactation, there is a need for coordination to compensate for prolactin (PRL) secretion, while during offspring weaning, this coordination is modified to halt PRL production and allow other hormones, such as follicle-stimulating hormone (FSH) and luteinizing hormone (LH), to reestablish the reproductive cycle. The values obtained during these phases are the lowest when compared to homotypic networks, which may more constrained pattern of signal coordination to meet hormonal output requirements. In the multipara condition, both networks reflects the lowest values, yet heterotypic interactions demonstrate an increased variance. This increase may be related to well-established integration in signal transmission, facilitating the anticipation of the next fertilization event and the subsequent initiation of hormonal cycling.

### 2.2 Signal classification and its relation to functional relevance

Recognizing the periodic and aperiodic differences across network interactions among different hormonal challenges, we next explored whether spontaneous activity could be grouped into signal classes. To achieve this, we employed a spectral decomposition algorithm for signal analysis [23]. Where under nullipara conditions, spontaneous activity could be categorized into two major classes, as illustrated in Figure 3A. This classification corresponds to the various conditions we evaluated. Notably, in most cases, the first two principal components from the spectral dimensionality reduction accounted for the majority of the variance in both networks, homotypic and heterotypic. This finding highlights the low dimensionality inherent in the pituitary system, which we will explore further in Subsection 2.5. Biologically, this suggests that the system is structured and conserved, with a few key variables effectively explaining its behavior. It also indicates that signal transmission is influenced by a limited range of frequencies or signal harmonics, which ties into the findings presented in Figure 2. It is important to highlight that signal classification is consistent in terms of wave shape. For example, Class 1 consists of cells that exhibit slower frequencies, while Class 2 primarily includes signals with higher frequencies (see Figure 3A). To explore the relationship between both classes and their aperiodic components, we measured the sensitivity of classes in relation to their aperiodic values. Our findings indicate that, in most instances, particularly in the homotypical network, the lowest aperiodic values (represented by cooler colors) were associated with cell signals from Class 1. Conversely, Class 2 was associated with higher aperiodic values, depicted by warmer colors, across various conditions and populations, as visualized through adjacency matrix arrangements (see Figure 3B). To validate this relationship, we employed surrogate resampling of aperiodic values from each population and condition to extract average values and internal variability. We observed that homogeneous populations consistently demonstrated a clear distinction between Class 1— characterized by lower mean aperiodic values — which tended to increase across the evaluated conditions in the homotypical population. In contrast, Class 2 showed an increase in mean aperiodic values, peaking notably in the weaned condition. This peak could be attributed to the necessity of halting hormone secretion, thus requiring greater inhibitory integration. In the case of the heterotypic population, there was no significant separation concerning Classes and aperiodic values, indicating reduced independence between them. This could be due to the influence of other endocrine cell groups contribute to the reshaping of network communication (Figure 3B, squared boxplot). Continuing our exploration of the differences between homotypic and heterotypic networks, our next question was focus on the amount of information (H) that both classes carry. However, the information distribution values did not indicate a preference between the two network types and remained relatively consistent across conditions (see Figure 3C, top panel). Notably, only the Class 2 (H) values during the weaned condition in the heterotypic network showed an amplification in rank values when compared to Class 1 of the same network, as well as Class 2 of the homotypic network. Building on our observations regarding the aperiodic value (see Figure 3B), we next sought a global relationship—without class distinctions—between these values. We found that the correlation values in homotypic networks were close to zero, negative, and non-significant, while the heterotypic networks exhibited a trend of increased anti-correlated values across conditions, showing significant correlations only in the lactanting, weaned, and multipara conditions (see Figure 3C, bottom panel). This suggests an inverse relationship between entropy values and aperiodicity; specifically, as the H entropy increases, the aperiodicity of the signal decreases under certain conditions.

**Figure 3:**
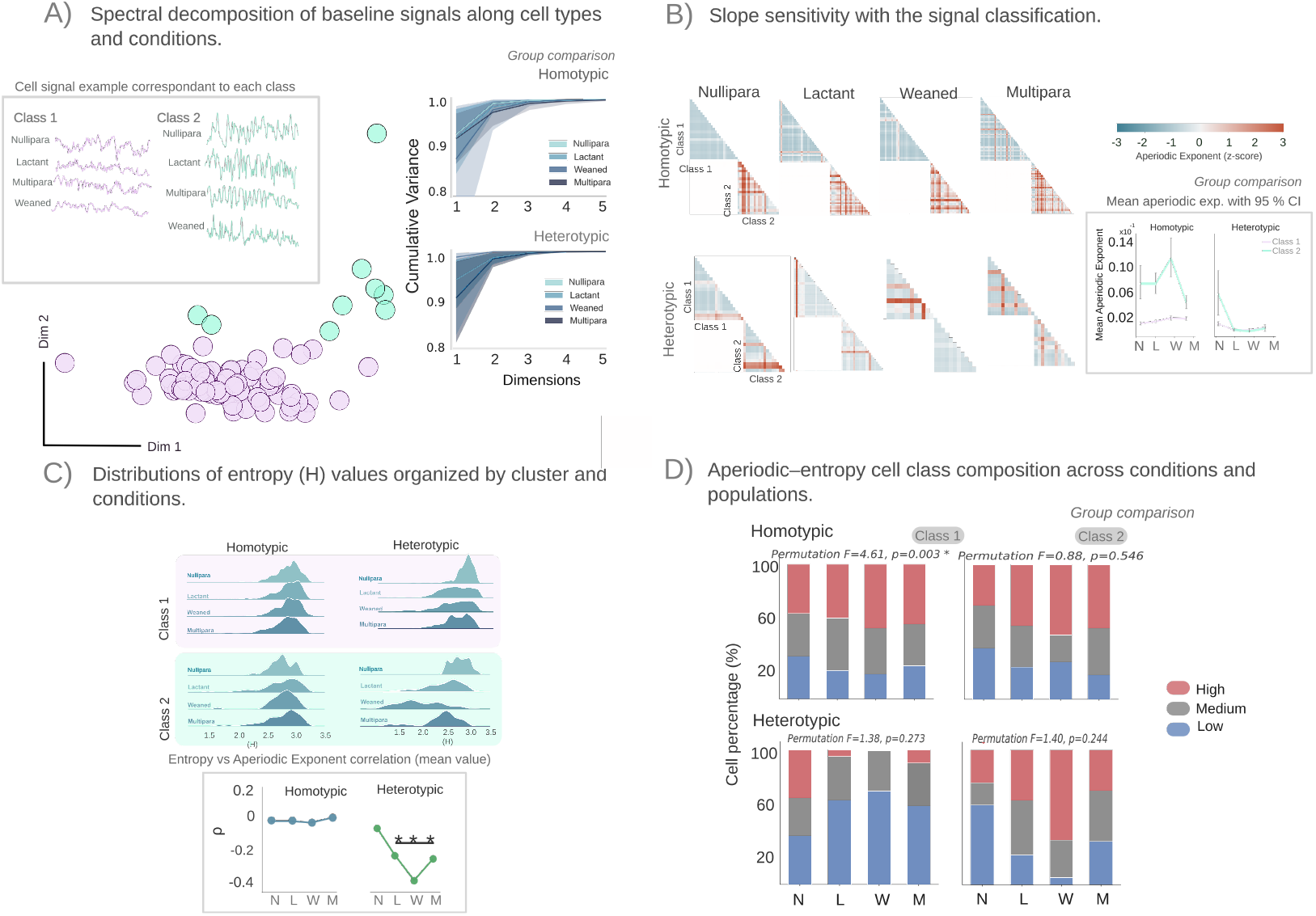
Sensitivity of the aperiodic exponent to spectral signal classification and entropy. **(A)** Left: spectral decomposition-based classification of spontaneous calcium time-series. Inset panels show representative examples of two signal classes: Class 1 (violet) and Class 2 (aqua). The scatter plot projects cell classes onto the first two principal dimensions (shown for multiparous lactotrophs). Top-right: cumulative variance explained across the five principal dimensions, for each cell population and condition (colour code as in Figure 1). **(B)** Group comparison of aperiodic slope as a function of signal class. The triangular matrix indicates the proportion of cells in Class 1 (upper triangle) or Class 2 (lower triangle), with the colourmap encoding the *z*-scored aperiodic exponent. Square panels display the mean aperiodic exponent ± 95% confidence intervals per class (colours as in panel A). **(C)** Group-level entropy (*H*) values across populations and conditions. Distributions are shown separately for Class 1 (top) and Class 2 (bottom), shaded in their respective class colours. Square panels show significant Spearman correlations (^∗∗∗^*p <* 0.05) between entropy and the aperiodic exponent, per population and condition. **(D)** Percentage of cells with low, medium, and high aperiodic-entropy co-classification values, stratified by oscillatory class (Class 1, right; Class 2, left) and population. Stacked bar charts show mean percentages across recordings (*n* = 4 per condition, individual values overlaid as points), with blue, gray, and red bars denoting low, medium, and high aperiodic—entropy bins, respectively, for homotypic (top) and heterotypic (bottom) populations across physiological conditions. Significant effect in Class 1 homotypic cells (*F* = 4.62, *p* = 0.003), with no individual pairwise comparison surviving correction. No significant effects were found in Class 1 heterotypic (*F* = 1.38, *p* = 0.27), Class 2 homotypic (*F* = 0.88, *p* = 0.55), or Class 2 heterotypic cells (*F* = 1.40, *p* = 0.24).

To explore this further, we quantified the proportion of cells exhibiting low, medium, and high aperiodic-entropy co-classification values within each oscillatory class, and assessed whether Class 1 or Class 2 showed greater sensitivity to physiological condition (Figure 3D). Differences in compositional distributions across conditions were evaluated using a multivariate permutation test applied to the joint [low, medium, high] percentage vector (pseudo-*F* statistic, *n*_perm_ = 10,000, Bonferroni correction for pairwise comparisons).

A significant overall effect of condition was detected in Class 1 homotypic cells (*F* = 4.62, *p* = 0.003), reflecting a shift away from the relatively balanced aperiodic-entropy composition observed in the nullipara condition toward a progressively higher proportion of high-aperiodic cells in lactant, weaned, and multipara animals. A qualitatively similar pattern was observed in Class 1 heterotypic cells, where the high-aperiodic fraction dropped markedly outside the nullipara condition (∼ 36% to *<*8%); however, this trend did not reach statistical significance (*F* = 1.38, *p* = 0.27), most likely due to the high inter-recording variability inherent to this mixed-cell population, whose composition is sensitive to the particular cellular ensemble sampled. No significant condition effects were detected in Class 2 cells for either population (homotypic: *F* = 0.88, *p* = 0.55; heterotypic: *F* = 1.40, *p* = 0.24).

Taken together, those indicates that the condition-driven reorganisation of aperiodic-entropy cell-state composition is specific to Class 1, and is most clearly expressed in homotypic lactotrophs. The selective sensitivity of Class 1 — but not Class 2 — to physiological state suggests that the two oscillatory classes are differentially coupled to the hormonal and metabolic demands of reproductive transition, with Class 1 cells undergoing a measurable redistribution toward higher aperiodic activity as lactation is established and maintained.

### 2.3 Signal synchronization and directionality across the time

Prior descriptions of network coordination in the pituitary gland [5, 8, 17] have relied predominantly on signal synchrony assessed through linear measures such as correlation. To contextualize our approach within this framework, we examined pairwise correlations across signals regardless of cell type, distinguishing positive (synchronous) from negative (asynchronous) values to assess whether their distributions differ between homotypic and heterotypic populations. Although significant correlations were observed, the values exhibited a roughly symmetrical distribution between positive and negative correlations across all evaluated conditions (Figure 4A). However, the proportion of positively versus negatively correlated cells differed markedly between the two populations. In the homotypical network, nulliparous conditions were characterized by a higher proportion of positively correlated cells and a correspondingly lower proportion of negatively correlated cells. As hormonal demand progressed, this pattern reversed: the proportion of positive correlations declined while that of negative correlations increased. In contrast, the heterotypical network exhibited near-equal proportions of positive and negative correlations under nulliparous conditions, followed by an increase in the positive correlation ratio during lactation; this ratio subsequently returned to near-nulliparous levels during weaning and in multiparous conditions. These divergent patterns suggest that the two network types encode and regulate signal communication through distinct mechanisms. Whereas the homotypical network displayed an inverse trend in correlation-ratio dynamics, converging toward near-equal proportions by the multiparous condition, the heterotypical network followed a more direct relationship with hormonal state, with the positive correlation ratio rising during periods of high hormonal demand and declining during weaning. Regarding the mean correlation values governing homotypical interactions (see Figure 4B), there is a trend of decreasing correlation values associated with reduced demands. However, in the multipara condition, there is a resurgence of positive values, which may indicate the sustained existence of interactions following the first fertilization. In contrast, heterotypical interactions start with low interval values during nullipara conditions, increase during lactation, but ultimately decline to even lower mean values than in the initial state. This difference may suggest a greater prevalence of long-term experience in homotypical networks compared to heterotypical ones. This is likely due to the specialization of the homotypical network [5], which does not necessitate greater cooperative integration of additional endocrine cell groups after multiple fertilizations.

**Figure 4:**
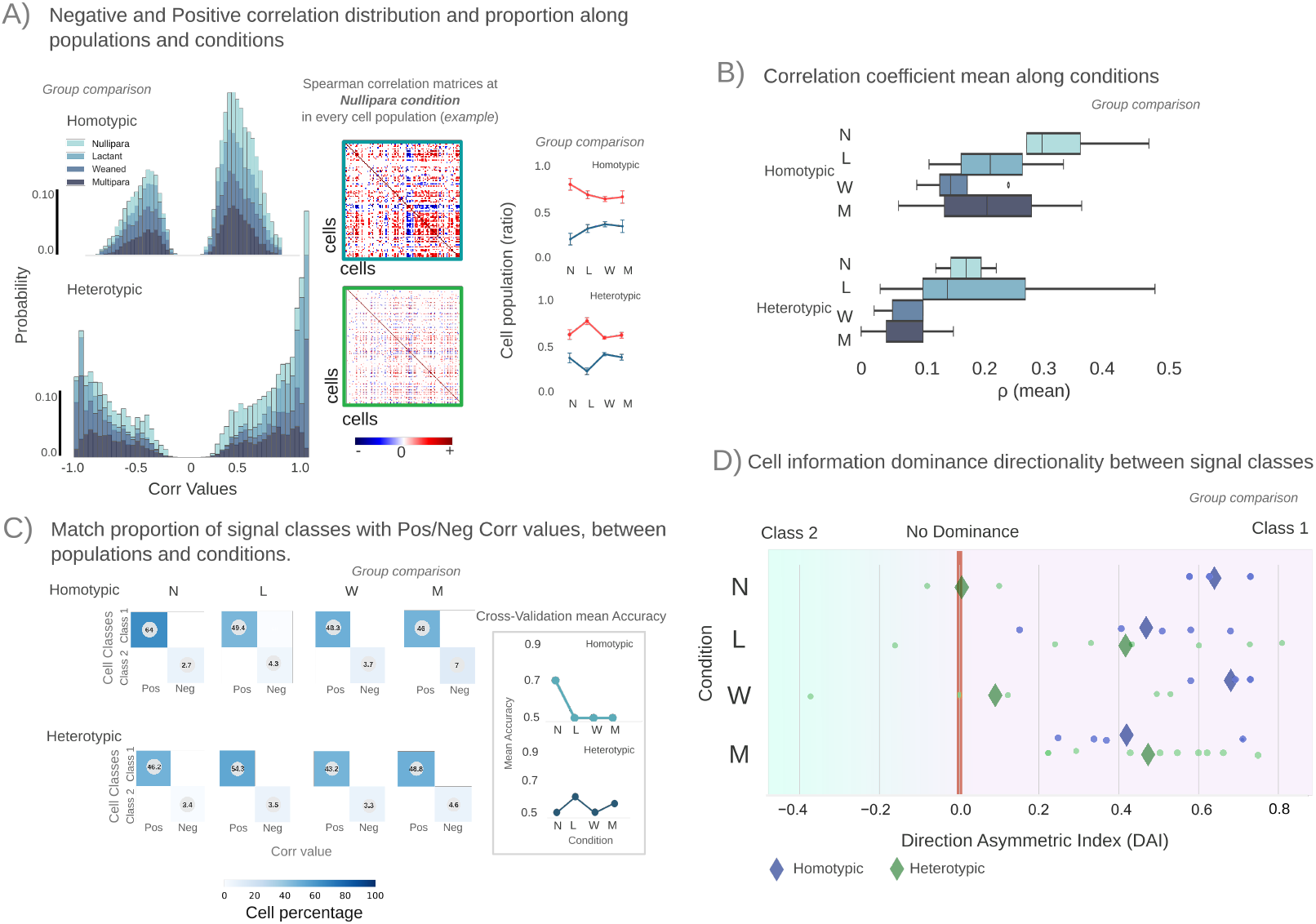
Synchronous and asynchronous dynamics, their relationship with signal classes, and directional information dominance. **(A)** Distribution of pairwise Spearman correlation coefficients across cell populations and conditions. The middle panel shows a representative adjacency matrix for the Nullipara condition, with distinct population clusters; positive correlations are shown in red, negative in blue. The right panel compares the proportion of cells displaying positive (synchronous) versus negative (asynchronous) correlations across conditions. **(B)** Mean Spearman correlation coefficient (*ρ*) across recordings and conditions. **(C)** Confusion matrices from random-forest classification, illustrating the relationship between signal class membership and correlation coefficient values. Entries and colour scale represent the percentage of true-positive cells. The left panel shows mean cross-validation accuracy, ordered as in the confusion matrices. **(D)** Direction Asymmetry Index (DAI) stratified by signal class, condition, and network type. DAI *<* 0 indicates Class 2 dominance; DAI = 0 indicates no dominance; DAI *>* 0 indicates Class 1 dominance. Individual sample values are shown as dots; the mean DAI per condition and population is indicated by a diamond symbol.

### 2.4 Dominance Directionality

Having broadly characterized the differences between the two network types, we next incorporated cell class into our classification approach to determine whether the correlation structure of the signals was also sensitive to class identity. After matching correlation values to their corresponding class and condition labels, we applied a random forest classifier to assess whether these variables could be used to discriminate between classes. The classifier performed poorly under most conditions, with the exception of the nulliparous homotypic condition (cross-validation accuracy > 0.5), where the largest separation between cell-class ratios was observed. Outside of this condition, cell class appeared largely unrelated to the connectivity inferred from the correlation metrics. Nonetheless, a consistent distinction between positive and negative correlation values was observed across both network types. Because correlation-based measures do not capture directionality, we next applied information-theoretic metrics to estimate directional interactions between signal classes (Figure 4D). In nearly all conditions, Class 1 was found to dominate signal communication; the sole exception was the nulliparous condition in heterotypic populations, in which no dominance between classes was observed. To further characterize this finding, we examined the dependence of class dominance on the embedding time of signals from both Class 1 and Class 2 (Figure 5). This analysis was motivated by the view that these results capture important features of network-level communication. As observed, both homotypic and heterotypic populations responded differentially to aperiodic activity, while showing no comparable sensitivity to information content or correlation values. Heterotypic populations additionally exhibited differences in cell integration related to the aperiodic exponent and to correlation values. Despite these distinctions, both population types displayed similar trends in directionality. Although the use of information theory allows the system to be approached probabilistically, this approach is not without limitations. In particular, our analysis relies on probability distributions derived deterministically from cell activity, which may limit the accuracy with which signal behavior can be predicted. To address this issue, we employed permutation tests and analyzed the lag axis to assess the direction of the signal (see to the Methods 10). Despite these challenges, analyzing how the system embeds under both homotypic and heterotypic network interactions allows us to preserve the geometric structure reflecting the shared evolution and synchronization of both classes over time. A representative example is shown in Figure 5A, where the embedding time space of Class 1 and Class 2 is projected onto the first two dimensions, with the heterotypic network shown at the bottom. Class 1 and Class 2 are visibly convergent at several time points (indicated by red dots), not only under nulliparous conditions but also during lactation, weaning, and the multipara stages. To further characterize this initial observation, we quantified the time points at which the two cell classes converged, focusing on the degree of synchronization between classes at different levels. We performed a Procrustes analysis on the embedding space of the two cell classes to determine the minimal distance between their trajectories. After aligning the trajectories, we measured local geometric synchronization across tiled windows (window size = 20) by factorizing the matrices derived from the embedding time series, thereby quantifying the degree of synchrony between classes. Values approaching 1 indicate substantial subspace overlap (strong synchrony), whereas values near 0 indicate orthogonal, uncorrelated dynamics. The resulting distributions of synchrony values are shown over time for each condition and population (Figure 5B). Notably, periods of low synchronization coincided with the emergence of dominance between trajectories, which we assessed using angular distances in the embedding space (Figure 5C). As shown in Figure 5B, dominance was evident across all conditions and both network types, as reflected by the principal angle separating the Class 1 and Class 2 trajectories; the distribution of angle values fell predominantly above the threshold expected under random alignment.

**Figure 5:**
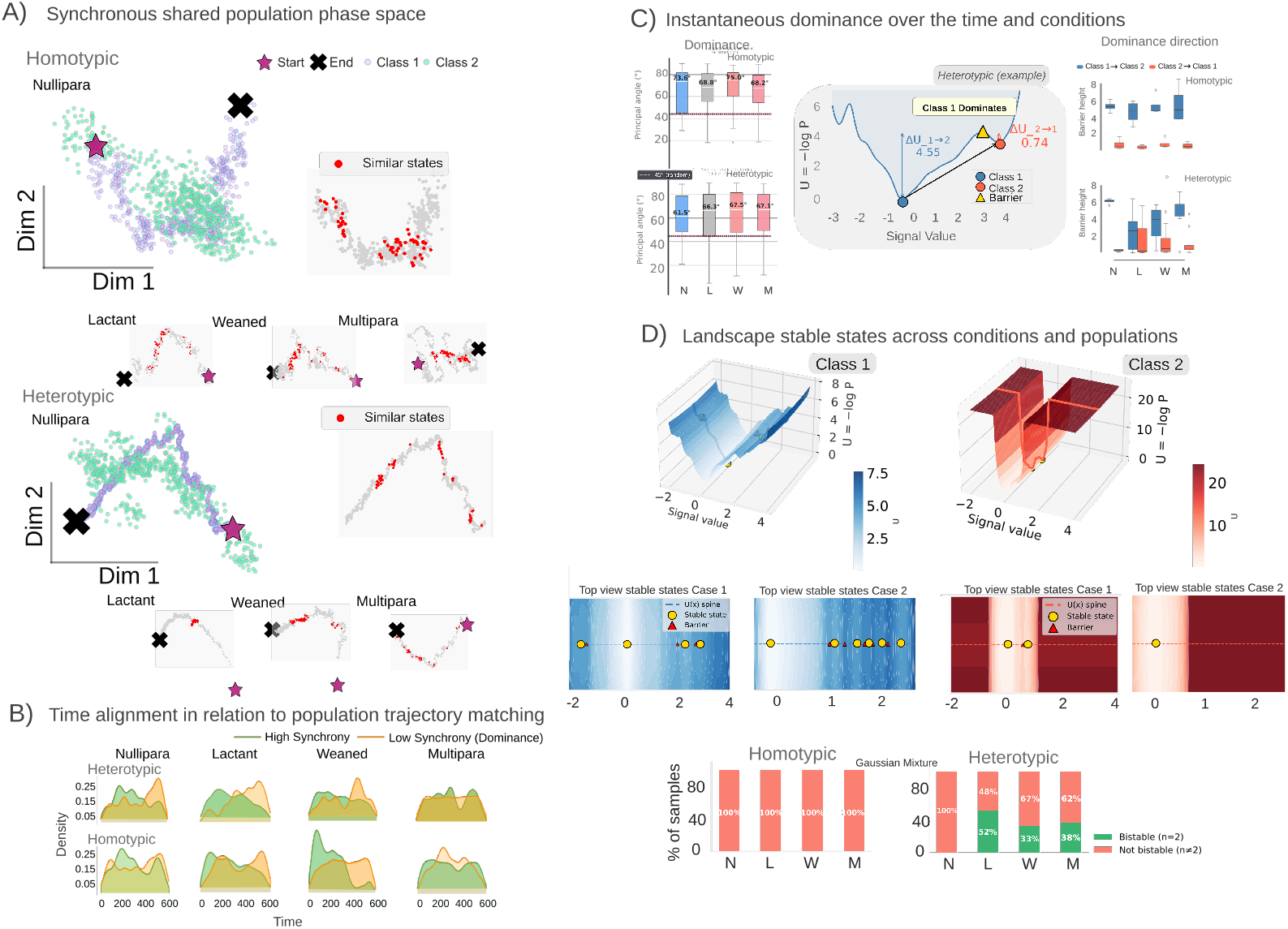
Geometric landscape structure of synchronous dynamical states. **(A)** Class synchrony in embedding time. Trajectories of two representative signal classes are shown for homotypic networks (top) and heterotypic networks (bottom). Grey shading marks the shared state-space regions visited by both trajectories, illustrating their subsequent divergence in representative examples from the Lactant, Weaned, and Multiparous conditions. **(B)** Temporal alignment relative to class trajectories. Green: probability density of trajectory points indicating high inter-class synchronisation. Orange: trajectory points exhibiting low synchronisation, associated with dominant-population activity. **(C)** Instantaneous dominance angles between Classes 1 and 2 across conditions and network types. Boxplots display principal angle values for significant dimensions, colour-coded by condition, for homotypic (top) and heterotypic (bottom) networks. The dashed line marks the chance angle of 45^◦^. The middle panel shows the spontaneous-activity quasi-potential landscape of a Nullipara heterotypic network, highlighting both fixed-point wells (Class 1 and Class 2) and the energy barrier separating them. Right boxplots depict barrier height between the Class 1 and Class 2 wells (arbitrary units). **(D)** Three-dimensional landscape representations for Class 1 (blue) and Class 2 (red), from a representative Weaned heterotypic network. Bottom: top-down views of Class 1 and two Class 2 examples illustrating a single stable fixed point. The bar chart shows the percentage of homotypic and heterotypic networks exhibiting bistability (more than one stable point) versus unimodality (single stable point), as determined by a Gaussian mixture test for multimodality.

To determine which class led the trajectory, we measured the height of the barrier separating the first peak associated with Class 1 from the subsequent peak associated with Class 2 (inset, Figure 5C), with a higher barrier indicating greater dominance. Across all homotypic conditions, Class 1 consistently occupied the dominant position (Figure 5C). In the heterotypic network, by contrast, dominance was more variable: Class 1 was dominant under nulliparous conditions, shifted toward shared dominance between classes during lactation and weaning, and appeared to return to Class 1 dominance in the multiparous condition. This variation points to a key functional distinction between the two network types. In the homotypic network, prior communication history and specialized cellular organization appear to support a stable, consistent response across varying hormonal demands. The heterotypic network, in contrast, appears to favor more cooperative dynamics during lactation and weaning, phases that likely require substantial reorganization, suggesting that dominance is more fluidly shared and exchanged between classes. This capacity for flexible dominance switching points to a broader principle: stability in both homotypic and heterotypic systems may not be a fixed property, but rather an emergent outcome of the interplay between signal dynamics and network composition.

### 2.5 Existence of momentaneous bistability in the spontaneous activity dynamics in both homotypic and heterotypic networks

Having observed that landscape dominance exhibited two well-defined points that differed in dominance between classes, we next investigated the stability of states within each class separately. The energy landscape for Class 1 exhibited one or more point-attractor states, although not all of these were stable. In contrast, the landscape for Class 2 was generally limited to one or two stable states, as illustrated in the middle panel of Figure 5D. These differences may account for the observed variation in dominance between populations and classes, since the presence of multiple stable points is thought to confer flexibility and robustness to a system [24].To evaluate this possibility, we performed a multistability test on the stable points identified in the landscape of each class (Figure 5D, bottom panel); because bistability was assessed jointly across both classes, the reported percentages of presence or absence reflect the combined behavior of Class 1 and Class 2. The presence of bistable or multistable dynamics in cell trajectories is suggestive of substantial underlying communication dynamics. In the homotypic network, most landscapes remained monostable, with only a small subset of samples exhibiting bistability, consistent with the unidirectional dominance observed in Class 1. The heterotypic network, by contrast, exhibited bistability during both lactation and weaning, with lactation showing the highest proportion of multistable landscapes, while the nulliparous condition showed no evidence of bistability. This pattern may further reflect the presence of cooperative, flexible dynamics within the heterotypic network. The relevance of bistable dynamics to the coordination of spontaneous secretory activity suggests that this system can dynamically switch between monostable and bistable regimes. This property is characteristic of non-linear biological systems that generate oscillatory behavior [24-27], and motivated our use of a generative model to investigate how geometric shifts can emerge from network-level coupling alone, without requiring external input. To determine whether the geometric coordination observed in the biological data could be recapitulated by such a minimal framework, we constructed a low-rank recurrent neural network (RNN) [28] composed of two coupled populations. Given the low-dimensional structure of the empirical trajectories (Figure 3A), each population was modeled as a leaky-integrator RNN with rank-2 recurrent connectivity. Network sizes were chosen to preserve the cellular disproportion observed in the biological system, with Population A comprising 80 units and Population B comprising 50 units (Figure 6A). The resulting synthetic time series qualitatively recapitulated the empirical data, with Population A dynamics resembling Class 1 activity patterns and Population B dynamics resembling those of Class 2 (Figure 6A, insets). Three coupling conditions were simulated to systematically probe the directionality of geometric coordination (Figure 6B). In Case 1 (directed condition), coupling was unidirectional from Population A to Population B, reflecting the hypothesized dominance of Population A observed in the biological data. In Case 2 (reversed condition), the direction of coupling was inverted, with Population B driving Population A, providing a critical test of whether geometric dominance follows the direction of network drive. In Case 3 (null condition), no coupling was present between populations, serving as a baseline against which the effects of directed coupling could be evaluated. Applying the same dimensionality-reduction pipeline used for the empirical data, Population A consistently occupied the dominant position in geometric space across all three conditions, coordinating trajectory structure independent of coupling direction (Figure 6B). This result was expected, as shifts in geometric dominance become apparent only once the subspace angle is explicitly quantified (Figure 6C), confirming that the generative model preserves the core geometric properties of the biological system. Having established the geometric validity of the model, we next examined how tonic external input modulates the latent dynamics under the directed coupling condition. Three input strengths were applied to Population A to emulate hypothalamic hormonal stimulation. In the absence of input, Population A exhibited concentric oscillations around the trajectory of Population B (Figure 6D). As input strength increased, the radius of these oscillations progressively decreased, with trajectories converging toward, but never reaching, a central attractor, a hallmark of a supercritical Hopf bifurcation [29] (Figure 6D–E). This behavior is mechanistically significant, as it indicates that the system sustains stable limit-cycle oscillations while remaining sensitive to input-driven amplitude modulation, thereby enabling transitions between slow and fast oscillatory regimes. Consistent with this interpretation, the temporal lag between Population A and Population B signals increased with input strength (Figure 6E), reflecting stronger entrainment of Population B by Population A under tonic stimulation. Together, these findings indicate that the intrinsic spontaneous behavior of the endocrine pituitary cell population, particularly that arising from heterotypic integration, supports transient bistable activity that enables transitions between slow and fast oscillatory regimes. Moreover, the predominance of slow oscillations, evident in both the geometric structure and the signal coupling, points to excitatory resonator-type dynamics within the system. This behavior remained consistent across conditions, including under baseline hormonal states and nulliparous conditions, underscoring the robustness of the system and its suitability for evaluation within a low-rank null model.

**Figure 6:**
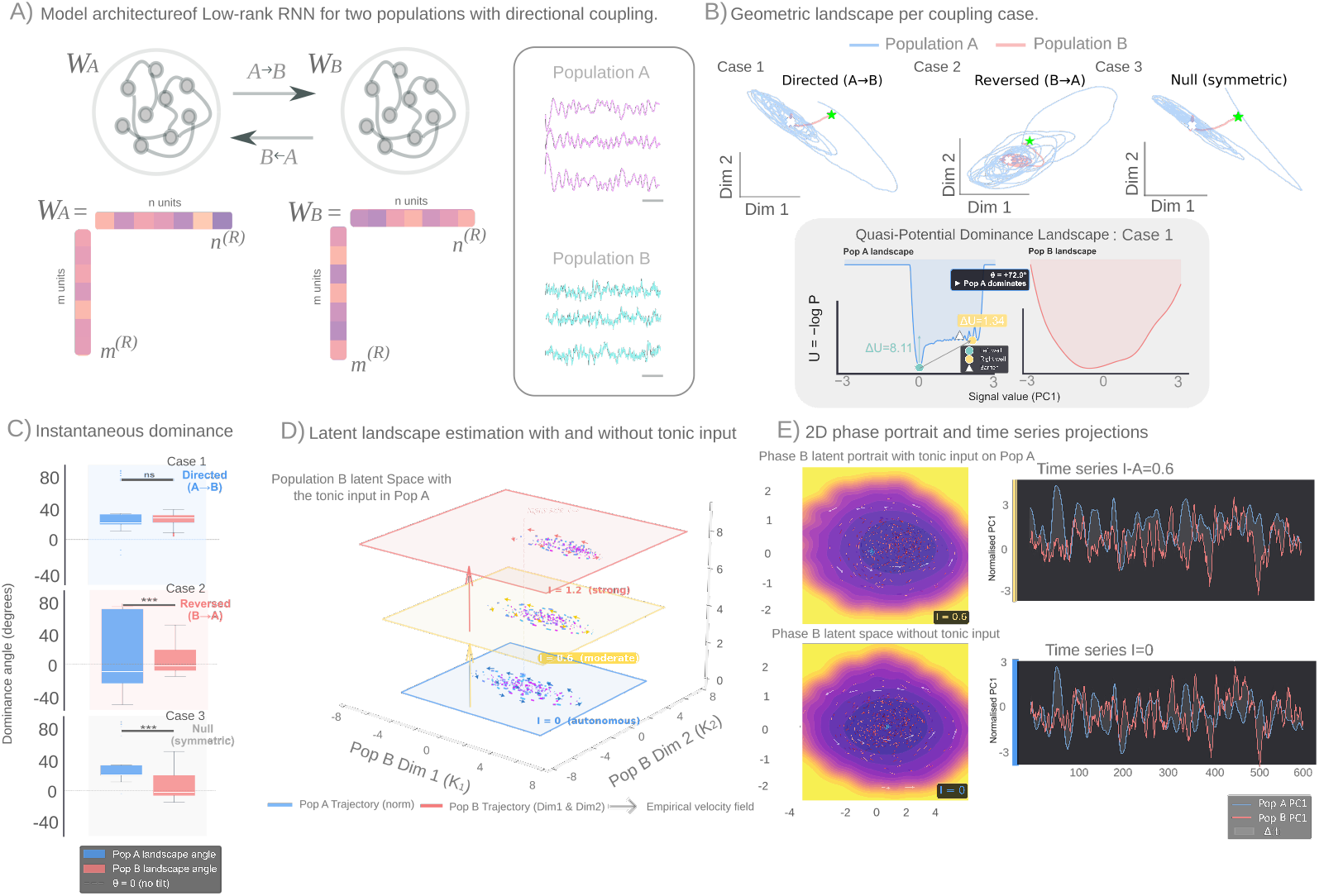
Low-rank dynamic geometric shift: generative null model. **(A)** Model architecture comprising two coupled rank-2 recurrent neural networks (*W*_*A*_ and *W*_*B*_) with asymmetric coupling strengths (*A* → *B* = 3.0, *B* → *A* = 3.0; null symmetric coupling = 0.0). Top: schematic of the coupling between population A (50 units) and population B (80 units). Bottom: construction of the weight matrix from the rank-1 vectors **m** and **n**. Left inset: three representative synthetic time series generated by the null model; the pink segment indicates epochs in which population A leads, and the blue segment indicates epochs in which population B leads. **(B)** Quasi-potential geometric landscapes for the three coupling cases. Case 1: directed connection *A* → *B* (star marks trajectory onset, cross marks end). Case 2: reversed direction *B* → *A*. Null case: symmetric coupling. The shaded bottom panel shows the quasi-potential landscape with the dominance angle of population A for Case 1. **(C)** Bootstrap instantaneous dominance angles from population A to population B. Angle *>* 0: population A dominates; angle *<* 0: population B dominates. Significance brackets indicate FDR-corrected Kruskal-Wallis tests per population (ns: non-significant; ^∗∗∗^*p <* 0.05). **(D)** Effect of tonic input (*I*) to population A on its latent representation within the population B latent space. Blue square: no input (autonomous oscillation); yellow: *I* = 0.6; red: *I* = 1.2. Magenta phase-portrait vectors correspond to population B; blue vectors to population A. Arrow direction and colour encode oscillation direction and input strength, respectively. **(E)** Two-dimensional phase portrait for Case 1, with and without tonic input. Background colourmap: energy landscape (magenta = high energy, yellow = low energy). Red trajectories: population B; blue trajectories: population A. White arrows indicate oscillation direction for *I* = 0.3 and *I* = 0. Right inset: representative time series projected onto the first principal component over 600 ms, illustrating how population A acts as a leader, increasing the temporal lag (Δ*t*) between the two populations when *I* = 0.6 (top example).

## 3 Discussion

The pituitary gland plays a crucial role in regulating hormone secretion. This process is fundamentally rooted in its ability to self-organize [1, 11, 30]. This autonomous structure enables the pituitary to maintain a coordinated output of hormones without needing external signals, all while supporting endocrine cells that can secrete hormones spontaneously [31, 32]. However, this spontaneity might also be organized by the emergence of structured oscillatory dynamics of intracellular calcium concentration [Ca^2+^]_*i*_ that are conserved across different pituitary cell types, regardless of their identity [3, 4]. Pituitary endocrine cells generate spontaneous action potentials and [Ca^2+^]_*i*_ transients driven by rhythmic Ca^2+^ entry [33]. In this study we addressed how these oscillations are classified, and how their coupling could establish a geometrical organization that governs spontaneous secretion at a system level. We combined data from homotypic lactotroph networks previously evaluated and published [5], with heterotypic interactions, allowing us to determine whether pituitary hormone secretion could be interpreted as reshaped across physiological states. In both network types, we evaluated spontaneous [Ca^2+^]_*i*_ dynamics using ex-vivo calcium imaging and extracted both, periodic and aperiodic signal components, where clear differences emerged in phase coupling and aperiodic exponent values depending on physiological stage (Figure 2), motivating a data-driven classification of oscillatory regimes (Figure 3). This revealed two functionally distinct cell classes: one characterized by slow oscillations (Class 1) and one by fast oscillations (Class 2). We then assessed their respective roles in network communication using both linear and nonlinear connectivity metrics (Figures 3–4), uncovering a consistent dominance direction originating from Class 1, alongside physiologically dependent differences in inferred connectivity strength and topology (Figure 4). Finally, we examined synchronization geometry in both classes using phase-space embedding methods (Figure 5), revealing differential dominance behavior between networks and a transient bistability that seeds a self-sustaining oscillator — one that maintains stable limit-cycle dynamics while remaining sensitive to amplitude modulation by external input (Figure 6). Together findings suggest that bistability in the oscillation dynamics between slow and fast cell types may represent a fundamental principle governing the function of the pituitary network. This property could allows the gland to adapt its secretory output in response to varying physiological demands. However, it also raises questions about how this intrinsic computational ability is altered during pathological conditions or prolonged hormonal stress.

### 3.1 Cell spontaneous classes intrinsically features

The observation of two functionally distinct oscillatory classes that modulate spontaneous intracellular calcium levels [Ca^2+^]_*i*_ is not new. Previous studies have described qualitative differences in oscillation waveforms, with particular emphasis on pseudo-plateau bursting. This pattern is characterized by sustained membrane depolarization, upon which bursts of action potentials are superimposed, resulting in high-amplitude [Ca^2+^]_*i*_ transients [13, 34]. For a long time, this bursting regime was thought to be the primary driver of hormone secretion, as larger fluctuations in calcium levels were linked to increased exocytotic output [13, 35]. Further ionic mechanisms have also been progressively characterized, including the blockade of BK channels in somatotrophs. This blockade converts bursting into continuous spiking and significantly reduces the amplitude of [Ca^2+^]_*i*_ oscillations [36]. Additionally, inhibiting voltage-gated calcium channels disrupts calcium influx, which is essential for sustaining both activity patterns [37]. What our data add to this picture is not simply the re-identification of these two regimes by our signal classification, but rather a network-level account of how they interact - and why their balance. Specifically, the dominance of one class alone does not appear sufficient to explain secretory coordination. Instead, our findings suggest that phase-coupled frequencies emerge within homotypic interactions under baseline hormonal conditions, indicating that network communication may already be functionally specialized even within populations of the same endocrine cell type. Nevertheless, it is important to acknowledge that purely homotypic interactions may represent an idealized constraint. In reality, there is known spatial intermixing of different endocrine cell types in the anterior pituitary. In fact, the close proximity of various cell types is likely the biological norm rather than the exception. In this context, the contribution of neighboring cell types to network organization is not only plausible but expected [10, 11, 38], and this is reflected in our data - the range of phase-coupled frequencies in heterotypic networks is broader than in homotypic ones, consistent with a more integrative communication architecture that is better suited to coordinate across oscillatory regimes. The aperiodic component of the signal offers a complementary window into these interactions. Building on work linking the 1/f signal spectral exponent to excitation/inhibition (E/I) balance, where a flatter power spectrum decay corresponds to a more excitable signal state and a steeper decay to a inhibition-domination. While these interpretations remain context and system-dependent, they provide a useful framework for approach to network cell organization [3–41]. Within homotypic networks, aperiodic exponent values showed a physiologically coherent pattern across conditions, Class 1 cells were associated with the lowest exponent values, consistent with a more excitable or functionally active state, while Class 2 cells exhibit the higher exponents, suggesting a relatively more constrained regime. Importantly, this class-specific aperiodic signature was corroborated by correlation synchrony values, which align with physiological expectations and provide independent mechanistic support for these observations. Heterotypic networks, in contrast, display distinct features in both, periodic and aperiodic domains, as well as in linear connectivity estimates, with class-specific aperiodic signatures becoming less pronounced. We propose this as a signature of cooperative signal integration, suggesting that cell types such as thyrotrophs and gonadotrophs could also be included in the network [42, 43]. In this expanded context, heterotypic integration may partially dissolve the strict class boundaries observed in homotypic settings, reshaping the overall communication topology in a physiologically meaningful way.

### 3.2 Bistability in geometrical coupling relevance

The emergence of a self-sustaining oscillator from the transient bistability observed between Class 1 and Class 2 populations is not simply a dynamical curiosity, it may represent a functional architecture with clear physiological implications. Limit cycle oscillators exhibit self-sustained oscillations, meaning they intrinsically persist in the absence of external periodic cues, and perturbation displacing the system from the cycle are damped out as the system asymptotically returns to its orbit [26, 27, 44]. The fact that the pituitary network operates in this regime — rather than as a noise-driven or input-dependent oscillator — has important consequences: it suggest that spontaneous hormone secretion is not merely a passive response to upstream signals, but an intrinsically generated output whose amplitude and timing can be modulated by input without depending on input for its existence.

This is precisely the logic of pituitary-adrenal ultradian rhythmicity, which until now has been proposed and explored only theoretically [45, 46]. What our phase-space Procrustes analysis adds is a geometric account of how this limit-cycle behavior is organized across populations. Rather than evaluating synchrony solely through pairwise signal correlations, we characterize the shape and alignment of the trajectories traced by each cell class traces through reconstructed attractor space. This approach is grounded in the neural manifold framework [47, 48], in which the joint activity of a population is represented as a trajectory in a high-dimensional state space, and the geometry of the same reflects the activity of the underlying network. Applied to the pituitary, this framework reveals that the two oscillatory classes do not merely differ in cell signal components, but also occupy distinct geometry roles within the collective dynamics of the system.

## 4 Limitations

The deep anatomical location of the pituitary gland, limits our ability to directly observe spontaneous population-level dynamics *in vivo*. In this study, we tackled this challenge by focusing on a mesoscale level of description. We used acute ex vivo pituitary slices, based on previously reported data [5] complemented by approaches that preserve local cellular architecture and intercellular communication pathways while allowing for experimental accessibility. Although this preparation provides a realistic framework for characterizing intra-pituitary cell dynamics, it also carries inherent limitation when interpreting our heterotypic network results. Because all spontaneously active cells were imaged collectively without prior cell-type identification or labeling, we cannot directly attribute the specific aperiodic or connectivity features of heterotypic interactions to defined endocrine populations such as thyrotrophs or gonadotrophs. Consequently, the cooperative restructuring of network communication described in the heterotypic condition was inferred from aggregate signal properties rather than from direct identification cell types. Future research work could address this limitation by using transgenic reporter lines, where specific pituitary cell populations express distinct fluorescent indicators under hormone-specific promoters. Combined with multi-laser excitation calcium imaging, this approach would allow simultaneous functional imaging of two or more endocrine cell types within the same preparation. Such an approach would directly test whether the class-specific dynamics and dominance relationships we describe follow cell-type boundaries, or instead reflect a more general organization principle independent of hormonal identity. Altogether, this methodology would significantly strengthen the hypothesis of cooperative heterotypic communication proposed in this study.

## 5 Conclusion

The findings presented in this study enhance our understanding of how the pituitary gland organizes spontaneous hormone secretion at a network level. Rather than attributing basal secretory activity to the intrinsic properties of individual cell types in isolation, we show that the collective dynamics of two oscillatory cell classes—characterized by distinct intracellular calcium ([Ca^2+^]_*i*_) signatures, aperiodic profiles, and phase-space geometry—form a functional architecture. This architecture exhibits an emergent property: a self-sustaining, bistability-seeded limit cycle. Through this mechanism, the anterior pituitary gland may integrate complex signaling by coordinating the interactions between different cell populations and signaling pathways, which is essential for maintaining physiological homeostasis. Our results provide a clear and robust dynamical mechanism supported by low-rank replicability—for this integrative capacity. The balance between slow and fast oscillatory regimes, together with the directed coupling that Class 1 cells exert over Class 2 cells, may enables the gland to sustain stable secretory rhythms while remaining responsive to physiological demands. This framework undeniably offers a compelling new perspective for understanding spontaneity in secretory disorders, including the disruption of secretory processes in hyper-functioning adenomas.

## 6 Resource availability

All analysis scripts and code required to reproduce the results presented in this study are publicly available at GitHub: https://github.com/AnaAquiles/GeoSync-Pituitary. Further information or resource will be send it under consideration. Raw datasets are available from the corresponding author upon reasonable request.

## 7 Acknowledgements

We gratefully acknowledge the valuable discussion on theoretical foundations with Pierre Fontanaud, as well as the insightful suggestions provided. The authors have received grants from the Agence Nationale de la Recherche (to P.M, ANR-22-CE14-0001), (to P.M, C.L, J.A.A, ANR-24-INBS-0005), D.J.H. was supported by MRC (MR/S025618/1), Diabetes UK (22/0006389) and UKRI ERC Frontier Research Guarantee (EP/X026833/1) Grants; T.F acknowledges support from CBF2023-2024-727 and IN206624. Finally A.A was supported by Foundation de la Recherche Medical grant (SPF202309017725).

## 8 Author contributions

A.A., M.P., F.T., and S.A.Y. contributed to the conception of the study and offered experimental method guidance. A.A.J and L.C performed the experimental procedure of calcium imaging recordings. A.A designed the algorithm, performed formal analysis and visualization, and drafted the manuscript. All authors contributed to the review and editing of the manuscript.

## 10 Methods

All procedures were conducted in accordance with the Animal Welfare guidelines of the European Community and were approved by the relevant institutional ethics committees.

### 10.1 Animals

Wild-type C57BL/6 female mice (2 month-old-wild-type) were studied across four reproductive states: nullipara ( *n* = 4), lactanting (8-10 days post-partum; *n* = 4), multiparous (second lactation, 8-10 days post-partum; *n* = 4), and weaned (dams weaned at postnatal day 15; *n* = 4). These conditions span the principal neuroendocrine states associated with prolactin and growth hormone secretion, allowing controlled comparison of pituitary network dynamics across reproductive demand.

### 10.2 Calcium Imaging Datasets

#### 10.2.1 Re-evaluated datasets

A subset of analyses was performed on previously published and publicly reported calcium imaging datasets [5]. For lactotroph populations, four physiological conditions were included: nullipara, lactanting (8-10 days post-partum), multiparous (second or third lactation), and weaned (females weaned at postnatal day 21). Full methodological details for these datasets are provided in the original publication [5].

#### 10.2.2 *In vitro* calcium imaging

Female mice at the reproductive stages described above were anaesthetised and decapitated. Pituitary glands were extracted and coronally sectioned into ∼200 µm slices using a vibratome; three slices per animal and condition were coated with poly-L-lysine, for imaging.

Slices were loaded with the calcium-sensitive fluorescent dye Fluo-4 AM (22 µM; Invitrogen, Eugene OR, USA) dissolved in dimethyl sulfoxide (DMSO; Sigma, St Louis MO, USA) with 0.02% pluronic acid F-127 (Sigma, St Louis MO, USA) in 1 mL of oxygenated Ringer solution (in mM: NaCl 138, KCl 2.5, NaH_2_ PO_4_ 1.25, NaHCO_3_ 3, HEPES 10, glucose 11, CaCl_2_ 2, MgCl_2_ 1). Dye loading was carried out for 45 min at 37C in a humidified atmosphere (5% CO_2_ / 5% O_2_).

Fluorescence imaging was performed on a fluorescence stereomicroscope (SteREO Discovery V12; Carl Zeiss, Oberkochen, Germany) equipped with a 20× water-immersion objective (WplanApo, 20Xw NA=1, Carl Zeiss), coupled to a scientific sCMOS camera (ORCA-Fire; Hamamatsu Photonics, Hamamatsu, Japan) operating at 4× pixel binning. Calcium signals were acquired at one frame per 200 ms (5 Hz) under constant perfusion with oxygenated Ringer solution at 34C for a total duration of 10 min per slice.

### 10.3 Time-Series Preprocessing

Raw fluorescence time series were preprocessed in four sequential steps to yield stable, comparable signals across cells and animals.

#### 10.3.1 Baseline normalisation

Raw fluorescence matrices were first transposed to the convention (cells × time). Each cell’s resting baseline *F*_0_ was defined as the minimum fluorescence value observed during the first 4,000 time samples, corresponding to an epoch prior to any stimulus-evoked activity. The full trace was then divided by *F*_0_ to produce a dimensionless ratio:

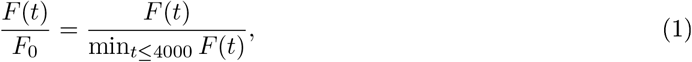

anchoring each cell’s signal to its own pre-stimulus baseline and correcting for inter-cell differences in absolute fluorescence intensity.

#### 10.3.2 Exponential drift correction

Slow multiplicative drift in the normalised signal – a characteristic artefact of photobleaching in calcium imaging – was removed by estimating and subtracting a shared exponential trend. A single linear regression was performed on the log-transformed *F/F*_0_ values pooled across all cells, yielding a global slope 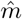 and intercept 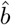 that characterise the dominant exponential decay:

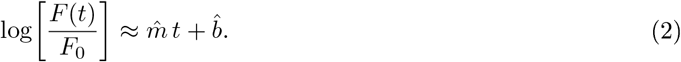

The fitted exponential 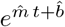 was then subtracted from each individual trace, producing drift-corrected signals with a stable baseline. Fitting a single trend to the pooled data ensures that the correction captures the component of variance shared across the population (i.e. global photobleaching) while preserving cell-specific dynamics.

#### 10.3.3 Low-pass filtering

Following drift correction, residual high-frequency noise was suppressed by applying a zero-phase Bessel low-pass filter (order 2, normalised cutoff frequency *f*_*c*_ = 0.3) along the time axis using forward-backward filtering (`scipy.signal.sosfiltfilt`). The Bessel design was chosen for its maximally flat group delay, which preserves the temporal waveform of calcium transients without introducing phase distortion [49].

### 10.4 Aperiodic Spectral Analysis

The aperiodic (1/*f*-like) component of the power spectrum was characterised for each cell by fitting a Lorentzian model to the single-sided power spectrum and extracting the aperiodic exponent. Cross-cell variability in log-power space was then summarised via frequency-resolved variance and mean profiles.

#### 10.4.1 Power spectrum estimation

For each cell *c*, the single-sided power spectrum was obtained from the discrete Fourier transform of the fluorescence trace *x*_*c*_(*t*) (*t* = 1, …, *N*_samples_):

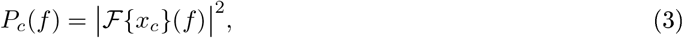

where ℱ{·} denotes the real-valued FFT and *f* ranges over the positive Nyquist grid with resolution Δ*f* = *f*_*s*_*/N*_samples_. Frequency bins at or below a threshold *f*_min_ = 0.5 Hz were discarded to exclude slow drift and DC components, yielding the trimmed spectrum 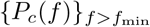.

#### 10.4.2 Lorentzian model of the aperiodic component

The aperiodic background of neural power spectra follows a 1/*f*^*α*^ power law that, in log-space, is well described by a Lorentzian function [40]:

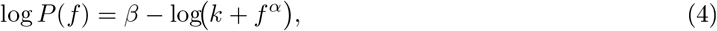

where *β* is a log-power intercept, *α >* 0 is the aperiodic exponent governing the spectral slope, and *k* ≥ 0 is a knee parameter that captures any low-frequency roll-off. In the limit *k* → 0 and for large *f*, Eq. (4) reduces to the pure power law log *P* (*f*) ≈ *β* − *α* log *f* .

#### 10.4.3 Aperiodic exponent estimation

The three parameters (*β, α, k*) were estimated for each cell by non-linear least squares minimisation of the residual sum of squares between the observed log-power and the Lorentzian model (Eq. (4)), using the Levenberg-Marquardt algorithm as implemented in `scipy.optimize.curve_fit` [49]. The aperiodic exponent was then defined as the absolute value of the fitted slope:

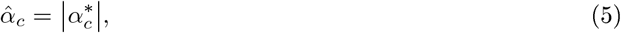

where 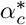 denotes the least-squares estimate for cell *c*. Taking the absolute value ensures that 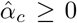 regardless of the sign convention returned by the optimiser, with larger values indicating steeper (more coloured) spectra.

#### 10.4.4 Cross-cell log-power variance

To quantify frequency-resolved heterogeneity across the cell population, the power spectra were transformed to log_10_-space with a small regularisation constant *ε* = 10^−10^ added to prevent numerical instability at bins with near-zero power:

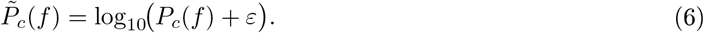

The cross-cell mean and variance were then computed at each frequency bin:

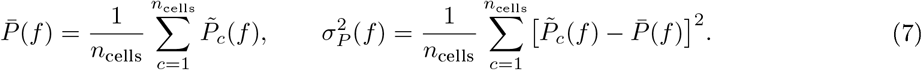

Frequency bins where 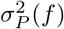 is large indicate spectral frequencies at which cells are most heterogeneous in their power, providing a data-driven criterion for identifying frequency bands of interest for subsequent analyses.

### 10.5 Phase-Amplitude Coupling Analysis

Phase-amplitude coupling (PAC) quantifies the degree to which the amplitude envelope of a high-frequency oscillation is modulated by the phase of a lower-frequency oscillation within the same signal. We computed PAC using the Modulation Index (MI) of [50], applied cell-by-cell to background-subtracted, exponentially normalised fluorescence traces (Δ*F/F, n*_cells_ cells).

#### 10.5.1 Frequency decomposition

For each candidate phase frequency *f*_*ϕ*_ and amplitude frequency *f*_*A*_, two narrow-band signals were obtained by applying a third-order Butterworth bandpass filter with passbands [*f*_*ϕ*_, *f*_*ϕ*_ + Δ*f*_*ϕ*_] and [*f*_*A*_, *f*_*A*_ +Δ*f*_*A*_], respectively, where Δ*f*_*ϕ*_ = 0.01 Hz and Δ*f*_*A*_ = 0.10 Hz. The instantaneous phase *ϕ*(*t*) and amplitude envelope *A*(*t*) were then extracted via the analytic signal obtained from the Hilbert transform:

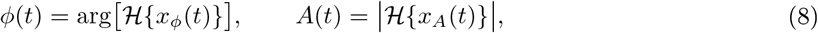

where *x*_*ϕ*_ and *x*_*A*_ denote the phase- and amplitude-filtered signals, respectively, and ℋ{·}is the Hilbert transform.

#### 10.5.2 Modulation Index

The full phase cycle [ −*π, π*) was partitioned into *N* = 18 equal bins. For each bin *k* the mean amplitude was computed as

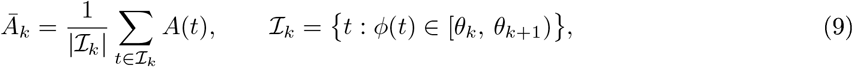

yielding a discrete amplitude-by-phase distribution 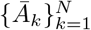. This distribution was normalised to a probability mass function

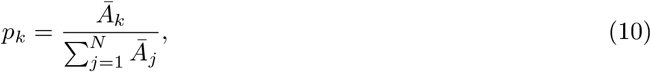

and its Shannon entropy was compared to the maximum entropy of a uniform distribution to obtain the MI:

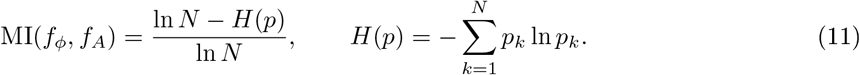

MI = 0 indicates a uniform amplitude distribution across phases (no coupling), whereas MI = 1 indicates complete concentration of amplitude at a single phase bin. A small regularisation constant (*ϵ* = 10^−12^) was added inside the logarithm to avoid numerical instability.

#### 10.5.3 Comodulogram

PAC was evaluated across a two-dimensional frequency grid. The phase frequency axis spanned *f*_*ϕ*_ ∈ [0.005, 0.045] Hz in steps of 0.005 Hz; the amplitude frequency axis spanned *f*_*A*_ ∈ [0.05, 0.75] Hz in steps of 0.05 Hz, yielding a 9 × 15 comodulogram per cell. The population-level comodulogram was obtained by averaging MI values across all cells:

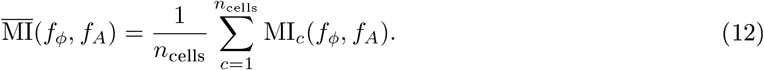

#### 10.5.4 Marginal profiles

To characterise how spectral power varies across cells as a joint function of frequency and time, principal component analysis (PCA) was applied to the per-cell spectrograms computed as described in Section

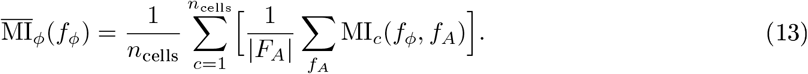

The dominant amplitude-frequency profile 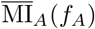 was computed analogously by collapsing the phase axis. Error bars in the corresponding figures represent ±1 SEM across cells.

### 10.6 Spectral Signal Classification

To characterise how spectral power varies across cells as a joint function of frequency *f* and time *t*, principal component analysis (PCA) was applied to the per-cell spectrograms computed as described before.

#### 10.6.1 Spectrogram covariance and PCA

For each cell *i*, the spectrogram *S*_*i*_(*f, t*) was evaluated over *F* = 60 logarithmically spaced frequency bins spanning 0.005-1 Hz and *T* non-overlapping time windows. The cross-cell covariance matrix was constructed over the joint (frequency × time) space as:

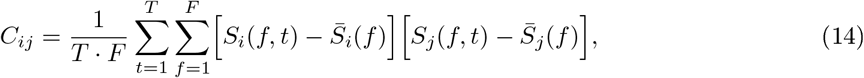

where 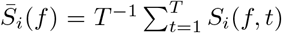 is the time-averaged spectrum of cell *i* at frequency *f*, and indices *i, j* run over all recorded cells. The maximum number of significant dimensions is bounded by *F* = 60, corresponding to the number of frequency bins.

The leading principal components - those capturing the largest fraction of total variance - were retained, and each cell’s spectrogram was projected onto these components to obtain a low-dimensional spectral embedding.

#### 10.6.2 Unsupervised clustering

Cells were grouped according to their spectral covariance structure by applying *k*-means clustering to the low-dimensional PCA projections, as implemented in,`scikit-learn` [51]. The number of clusters was set to *k* = 2, with *n*_init_ = 10 independent initialisations to mitigate sensitivity to centroid initialisation; the solution minimising the within-cluster sum of squares was selected. The cluster bandwidth was estimated directly from the data prior to fitting.

### 10.7 Aperiodic Signal Decomposition and Cluster-Level Analysis

To examine the relationship between spectral cluster membership and aperiodic signal properties at single-cell resolution, a cell-by-cell adjacency matrix was constructed for each combination of cell population (heterotypic and homotypic) and physiological condition (nullipara, lactanting, weaned, and multiparous). Spectral class indices and per-cell aperiodic exponents were merged at the level of individual cells using the recording index and cell identifier as joint keys; only cells present in both datasets were retained for subsequent analysis.

#### 10.7.1 Cluster size balancing

To prevent asymmetries in matrix structure driven by cluster size imbalance, clusters were downsampled to a common size prior to matrix construction. The minimum cluster size across all clusters within a given condition was determined, and any larger cluster was randomly subsampled without replacement to match this reference size (random seed fixed at 42 for reproducibility). A condition was included in the analysis only if both clusters contained at least one cell after merging.

#### 10.7.2 Aperiodic adjacency matrix

For each condition, an *N* × *N* symmetric adjacency matrix **A** was constructed, where *N* is the total number of retained cells across both clusters. Diagonal entries encode each cell’s intrinsic aperiodic exponent:

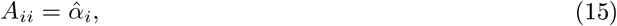

where 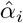 is the aperiodic exponent of cell *i* estimated as described before. Off-diagonal entries represent a pairwise aperiodic similarity measure defined as the arithmetic mean of the exponents of cells *i* and *j*:

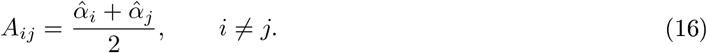

All matrix entries were subsequently *z*-score normalised across the full set of elements to centre the distribution and express values in units of standard deviation from the population mean:

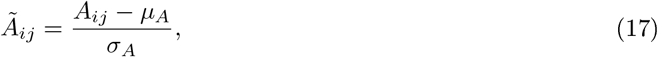

where *µ*_*A*_ and *σ*_*A*_ are the mean and standard deviation computed over all entries of **A**. The lower triangular portion of **Ã** was visualised as a heatmap using a diverging colormap centred at zero, with rows and columns ordered by cluster identity to allow visual inspection of within-versus between-cluster aperiodic structure. Cluster boundaries were annotated along both axes.

#### 10.7.3 Bootstrap Estimation of Within-Cluster Aperiodic Reliability

To assess the reliability of the mean aperiodic exponent within each spectral cluster, a non-parametric bootstrap procedure. For each combination of cell population, physiological condition, and cluster identity, the observed aperiodic values 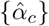 were resampled with replacement over *B* = 1,000 iterations. The mean aperiodic exponent was computed for each bootstrap replicate *b*:

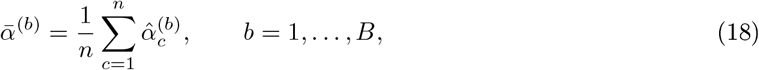

where *n* is the number of cells in the cluster and 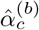 denotes the *c*-th draw of the *b*-th replicate. This procedure yields an empirical sampling distribution of the mean from which the 95% confidence interval was derived as the interval between the 2.5th and 97.5th percentiles:

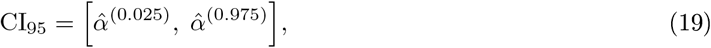

where 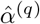 denotes the *q*-th quantile of the bootstrap distribution. Results were aggregated across all populations, conditions, and clusters; the observed mean together with its bootstrap confidence interval was retained as the summary statistic for each group.

### 10.8 Contingency Analysis of Aperiodic Exponent and Shannon Entropy

To examine the joint distribution of aperiodic signal properties and spectral entropy across cluster identities, both continuous measures were discretised and analysed via cross-tabulation at three levels of granularity.

#### 10.8.1 Tertile discretisation

The aperiodic exponent 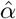 and the Shannon entropy *H* were each independently binned into population tertiles, partitioning the observed range into three ordered categories - Low, Medium, and High - each containing approximately one-third of the observations:

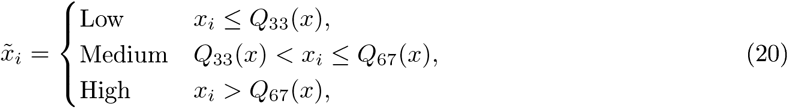

where *Q*_33_ and *Q*_67_ denote the 33rd and 67th sample percentiles, respectively. Tertile boundaries were computed independently for 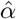 and *H*, so no assumption of a shared distribution is imposed.

#### 10.8.2 Cross-tabulation

Contingency tables were constructed at three levels of aggregation: **Pooled**: all cell populations and physiological conditions combined, providing a global view of the 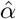-*H*-cluster relationship.**Population-level**: a single cell population (e.g. homotypic or heterotypic) with all conditions pooled.**Group-level**: a single cell population within a single physiological condition (e.g. lactotrophs during lactation). At each level, three tables were derived: (i) the marginal cross-tabulation of 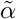 against cluster identity, (ii) the marginal cross-tabulation of 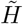 against cluster identity, and (iii) the joint cross-tabulation of 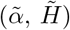 against cluster identity.

#### 10.8.3 Percentage representation

The joint contingency table was further expressed as a percentage of the total number of cells *N* in the corresponding subset:

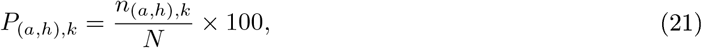

where *n*_(*a,h*),*k*_ is the count of cells falling in tertile category 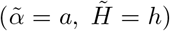 and belonging to cluster *k*, and *N* = ∑_*a,h,k*_ *n*_(*a,h*),*k*_ is the total cell count in the subset. This normalisation allows direct comparison of cell proportions across groups of different sizes and was used as the primary input for figure generation.

#### 10.8.4 Comparison of compositional distributions across conditions

To assess whether the joint aperiodic-entropy cell-state composition differed across physiological conditions, a multivariate permutation test was applied separately for each oscillatory class (Class 1 and Class 2) and each cell population (homotypic and heterotypic). For each recording *r*, the percentage values *P*_(*a,h*),*k*_ were collapsed across entropy position by summing within each aperiodic tertile, yielding a three-component compositional vector **v**_*r*_ = [*p*_Low_, *p*_Medium_, *p*_High_ ]_*r*_ that sums to 100% per recording. This compositional structure — whereby the three components are mutually exclusive and exhaustive - precluded the use of standard univariate tests applied independently per bin, as such an approach would violate the sum constraint and introduce spurious correlations between test statistics.

A pseudo-*F* statistic was used as the test statistic, defined as the ratio of between-group to within-group dispersion in compositional space:

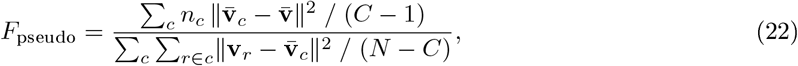

where *C* is the number of conditions, *n*_*c*_ is the number of recordings in condition *c*, 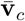 is the centroid of condition *c* in compositional space, 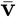 is the grand centroid, and *N* is the total number of recordings. The null distribution of *F*_pseudo_ was obtained by randomly permuting condition labels across recordings and recomputing the statistic over *n*_perm_ = 10,000 iterations. The permutation *p*-value was defined as the proportion of permuted statistics exceeding or equal to the observed value.

Following the omnibus test, pairwise permutation tests were conducted for all condition pairs using the same pseudo-*F* statistic restricted to the two groups under comparison. Resulting *p*-values were adjusted for multiple comparisons using the Bonferroni method (*m* = 6 pairs for four conditions).

### 10.9 Surrogate-Thresholded Pairwise Correlation Analysis

To identify statistically significant pairwise interactions among cells, a Fourier phase-randomisation surrogate procedure was applied to define a cell-pair-specific significance threshold for the observed Spearman correlation matrix.

#### 10.9.1 Fourier surrogate generation

For each cell *i*, the discrete Fourier transform of the fluorescence trace was computed and decomposed into its amplitude and phase spectra:

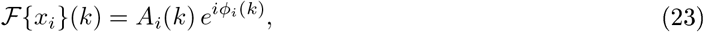

where *A*_*i*_(*k*) = |ℱ{ *x*_*i*_} (*k*)| and *ϕ*_*i*_(*k*) = arg[ℱ{*x*_*i*_*}* (*k*)]. For each of *B* = 1,000 surrogate resamples, the phase spectrum of every cell was independently replaced by phases 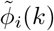 drawn uniformly from [− *π, π*), while the amplitude spectrum was held fixed. Conjugate symmetry was enforced to ensure a real-valued inverse transform:

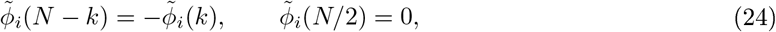

where *N* denotes the number of time points. The surrogate signal was then obtained as

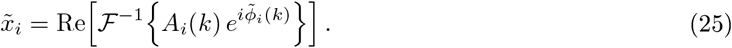

This procedure preserves the power spectrum of each individual cell while destroying inter-cell phase relationships, thereby generating a null distribution of pairwise correlations under the hypothesis of no genuine synchrony.

#### 10.9.2 Significance threshold and matrix thresholding

For each cell pair (*i, j*), the element-wise mean *µ*_*ij*_ and standard deviation *σ*_*ij*_ were computed across the *B* = 1,000 surrogate Spearman correlation matrices. An observed correlation *ρ*_*ij*_ was deemed significant only if it exceeded the surrogate threshold:

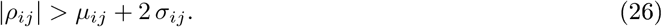

Correlations that did not satisfy this criterion were set to zero, yielding a sparse, thresholded correlation matrix 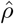 that retains only interactions whose magnitude is unlikely under the phase-randomisation null.

#### 10.9.3 Interaction classification

Non-zero entries of 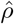 were partitioned into three interaction classes based on their magnitude and sign relative to the surrogate distribution: **Strongly synchronous**: 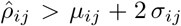. **Negatively asynchronous**: 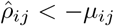 and 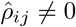.**Wea ly asynchronous**: 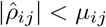 and 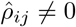.

The proportion of positive and negative interactions among all surviving (non-zero) pairs was computed, and the distribution of values within each class was visualised as a boxplot.

### 10.10 Cluster-Synchrony Association via Random Forest Classification

To assess whether spectral cluster identity is predictive of pairwise synchrony class, a supervised classification analysis was performed. Spectral cluster labels and synchrony labels were merged at the cell level using the recording index, cell identifier, and physiological condition as joint keys; only cells present in both tables were retained.

Synchrony labels were encoded numerically (synchronous = 1, asynchronous = 2). Cluster identity was used as the sole predictor feature and synchrony class as the target variable. A Random Forest classifier (implemented in `scikit-learn` [51] ; random seed fixed at 42) was evaluated by stratified 5-fold cross-validation, with classification accuracy as the performance metric. The mean and standard deviation of accuracy across folds are reported as the summary statistic. A confusion matrix was constructed to visualise the correspondence between cluster identity and synchrony class for each population-condition combination.

### 10.11 Transfer entropy along cell clusters

This methodology section was inspired on the previous reported in Aquiles 2025 [52], but further described here. We computed the transfer entropy from baseline in every cluster, ie *X*_*t*_ = signal from cellclass*n* and *Y*_*t*_ = signal from cellclass *m*; using a finite set of lags (100, (-50,50))) *L* = {*τ*_1_, *τ*_2_, …, *τ*_*k*_*}*. For each lag, we define the lagged version of the time series:

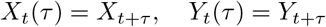

Then we calculate the Mutual Information (MI) between *X*_*t*_(*τ*) and *Y*_*t*_ as:

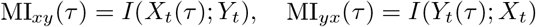

Where *I* corresponds to Shannon mutual information, and *H* to Shannon entropy:

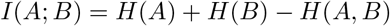

Where *A* and *B* represent the different cluster sets. To compute directionality, we took the vector MI_*xy*_(*τ*) for all *τ* ∈ *L*, where we consider the best delay as the one that maximizes the mutual information:

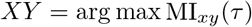

And we follow the same logic in a similar way for the other direction *Y X*.

#### 10.11.1 Multi-Set Directionality Computation

The directionality analysis is extended to multiple signal sets by evaluating all pairwise interactions among them. Each pair of sets is analyzed independently following the same time-lagged MI and permutation-testing procedure, see below.

#### 10.11.2 Directionality Asymmetry Index

To summarize net directional influence between two signal sets, the Directionality Asymmetry Index (DAI) is computed. Total significant information transfer is summed separately for each direction, and the DAI is defined as

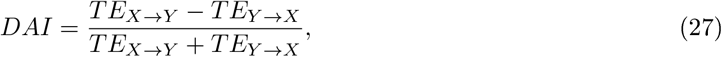

where *TE*_*X*→*Y*_ and *TE*_*Y* →*X*_ denote the total information transfer in each direction. The DAI ranges from −1 to 1, indicating dominant directional influence or balanced interactions.

#### 10.11.3 Permutation-Based Significance Testing of Mutual Information

To determine whether the observed mutual information (MI) at a given lag exceeded chance levels, a permutation null distribution was constructed independently for each lag *τ* . Specifically, *N* = 1,000 surrogate time series 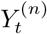 were generated by randomly shuffling the original *Y*_*t*_ (and equivalently *X*_*t*_ for the reverse direction). The MI was recomputed for each surrogate, yielding a null distribution. Only lag values for which the observed MI exceeded the 95^th^ percentile of the null distribution (*p <* 0.05, one-tailed) were retained for subsequent directionality analyses.

### 10.12 Geometric Synchrony Analysis Between Neural Populations

To quantify the degree to which two pituitary cell populations (PopA and PopB) share geometric structure in their collective dynamics, a manifold-based synchrony analysis was performed. The pipeline comprises five sequential steps: time-delay embedding, dimensionality reduction, Procrustes alignment, and two complementary synchrony metrics evaluated at global and local temporal scales.

#### 10.12.1 Time-delay embedding

Each cell’s fluorescence trace *x*_*i*_(*t*) was embedded into a pseudo-phase-space attractor using the Takens time-delay construction:

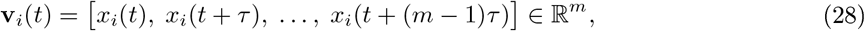

where *m* = 2 is the embedding dimension and *τ* = 1 sample is the time delay. The embedded trajectories of all cells within a population were concatenated horizontally, yielding a population-level state matrix 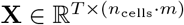, where *T* is the number of embedded time points.

#### 10.12.2 Dimensionality reduction via PCA

Principal component analysis was applied independently to the state matrix of each population, retaining up to *k* = min(10, *n*_cells_ · *m*) components:

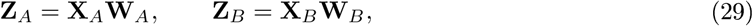

where **W**_*A*_ and **W**_*B*_ are the matrices of leading eigenvectors of the respective sample covariance matrices. This yields low-dimensional latent trajectories **Z**_*A*_, **Z**_*B*_ ∈ ℝ^*T* ×*k*^ that capture the dominant variance structure of each population in its own coordinate frame.

#### 10.12.3 Procrustes alignment

Because the PCA coordinate frames of PopA and PopB are independently defined, a direct geometric comparison requires a common reference frame. An optimal orthogonal rotation **R** was found by solving the orthogonal Procrustes problem:

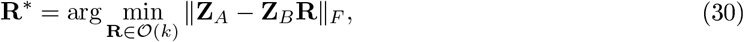

where O(*k*) denotes the group of *k* × *k* orthogonal matrices and ∥ · ∥_*F*_ is the Frobenius norm. The solution is given by **R**^∗^ = **VU**^⊤^, where **UΣV**^⊤^ is the singular value decomposition of 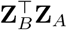. The aligned trajectory 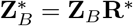 then lives in the same coordinate frame as **Z**_*A*_, enabling direct pointwise comparison.

#### 10.12.4 Geometric synchrony metrics

##### Pointwise trajectory distance

The instantaneous divergence between the two population states was quantified by the Euclidean distance between their aligned latent representations at each time point:

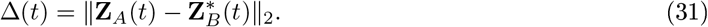

Time points where Δ(*t*) falls below its temporal mean are interpreted as epochs of high geometric synchrony; time points above the mean indicate divergent (asynchronous) population states.

##### Global principal subspace angles

To characterise the overall geometric relationship between the two population subspaces, the principal angles *θ*_1_ ≤ · · · ≤ *θ*_*k*_ between the column spaces of **Z**_*A*_ and 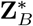 were computed via QR decomposition followed by SVD of the cross-product of the orthonormal bases:

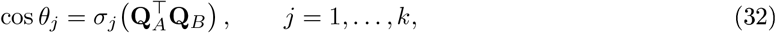

where **Q**_*A*_ and **Q**_*B*_ are the orthonormal factors from the QR decompositions of **Z**_*A*_ and 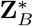, respectively, and *σ*_*j*_(·) denotes the *j*-th singular value. Angles below 45° indicate shared geometric structure (synchrony); angles exceeding 45° indicate geometrically divergent subspaces (dominance).

##### Sliding-window subspace alignment score

To capture the temporal evolution of geometric synchrony, a sliding-window variant of the principal angle analysis was applied. At each time step *t*, local segments of half-width *w* = 20 samples were extracted from both trajectories, and the mean cosine of the local principal angles was computed:

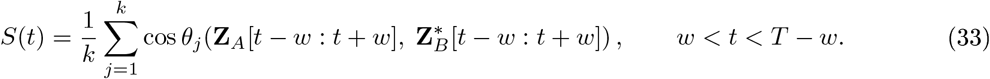

*S*(*t*) ranges from 0 (orthogonal, maximal divergence) to 1 (identical subspaces, perfect alignment). Values above the temporal mean of *S* are classified as high-alignment (synchronous) epochs; values below the mean as low-alignment (dominant) epochs.

#### 10.12.5 Temporal density estimation of synchrony epochs

To identify at which periods of the recording synchrony and dominance preferentially occur, a kernel density estimate (KDE) with a Gaussian kernel (bandwidth *h* = 0.2, Scott’s rule) was fitted separately to the set of time indices classified as high-alignment (top quartile of *S*(*t*)) and low-alignment (bottom quartile of *S*(*t*)). The resulting density curves provide a continuous representation of the temporal concentration of each synchrony regime across the recording.

### 10.13 One-Dimensional Quasi-Potential Landscape and Dominance Angle

To quantify the stability and relative dominance of two coexisting population states, the quasi-potential energy landscape was estimated from the empirical signal distribution of each cell group and physiological condition. This analysis operates on a one-dimensional projection of the population signal - complementing the two-dimensional latent-space landscape described before — and provides an explicit scalar measure of bistability and directional dominance.

#### 10.13.1 Quasi-potential estimation

For each scalar signal distribution 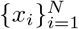, the probability density *P* (*x*) was estimated by a Gaussian kernel density estimator (KDE):

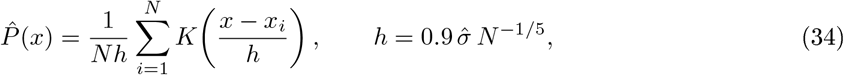

where *K* is the standard Gaussian kernel and 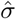 is the sample standard deviation. The quasi-potential was then defined as the negative log-density, regularised to avoid numerical instability at grid points with near-zero probability:

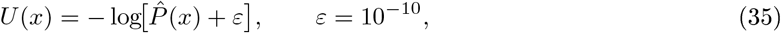

evaluated on a uniform grid of 500 points spanning the observed signal range. The landscape was shifted so that its global minimum equals zero: *U* ← *U* −min(*U*). Minima of *U* correspond to preferred (high-probability) population states; the value of *U* at each minimum is proportional to the negative log-probability of that state.

#### 10.13.2 Well and barrier identification

Local minima and maxima of *U* were identified using a neighbourhood-based extremum detector with order parameter *q* = 15 grid points (`scipy.signal.argrelmin` and `argrelmax`). A bistable landscape was defined as one containing at least two local minima separated by at least one local maximum. When multiple minima were present, the two deepest (lowest *U*) were retained as the primary wells; the highest local maximum strictly between these two wells was taken as the energy barrier.

Denoting the left and right wells as *A* and *B*:

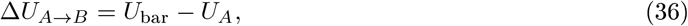

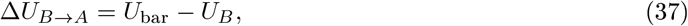

where *U*_ba_ is the barrier height. The mean escape time from well *P* is proportional to 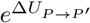, so a larger barrier corresponds to a longer-lived (more stable) state.

#### 10.13.3 Dominance angle

The relative stability of the two wells was summarised by the dominance angle *θ*, defined as the signed inclination of the line connecting well *A* to well *B* in the (*x, U*) plane:

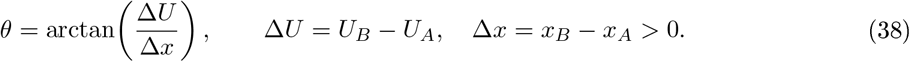

Because well *A* is always to the left of well *B* by construction (Δ*x >* 0), the sign of *θ* is determined entirely by Δ*U* :

- *θ >* 0: well *B* is higher than well *A*; population *A* occupies the deeper well and is the dominant state.
- *θ <* 0: well *A* is higher than well *B*; population *B* dominates.
- *θ* ≈ 0: symmetric landscape; neither population is strongly preferred.

The dominant population is formally defined as the one occupying the well with the lower *U* value (higher probability density). The magnitude |*θ*| quantifies the degree of asymmetry: larger angles correspond to stronger dominance and greater stability of the preferred state.

The dominance angle analysis was applied to three signal distributions per recording: the pooled signal across both populations, and the signals of each group separately. Results were aggregated across all physiological conditions (nullipara, lactanting, weaned, multiparous) and all recordings, yielding a condition-level summary table of *θ*, Δ*U*, and the dominant population for each group.

### 10.14 Bistability Analysis: Mixture Model Selection and State Dwell Times

To characterise the degree of bistability in pituitary endocrine cell populations, the empirical distribution of fluorescence signals was analysed using Gaussian Mixture Model (GMM) selection and per-cell dwell-time statistics. All analyses were applied per recording (sample) and condition, with results aggregated across samples for reporting.

#### 10.14.1 Signal pooling and quasi-potential estimation

For each recording, fluorescence traces were flattened across cells and time to yield three one-dimensional signal distributions: (i) all cells pooled (*X*_all_), (ii) Class 1 cells only (*X*_1_), and (iii) Class 2 cells only (*X*_2_). The quasi-potential landscape of each distribution was estimated as described in Section, Eq. (35), using a Gaussian KDE with Silverman’s bandwidth on a uniform grid of 500 points spanning the observed signal range. For multi-sample conditions, individual-sample landscapes were interpolated onto a common grid and averaged to obtain a condition-level mean landscape with pointwise standard deviation shown as a shaded band.

#### 10.14.2 Gaussian Mixture Model selection

To test whether the pooled signal distribution *X*_all_ was consistent with a bimodal (bistable) regime, Gaussian Mixture Models with *n* ∈ {1, 2, 3} components were fitted by Expectation-Maximisation. Each model was initialised *n*_init_ = 10 times with random starting parameters (random seed fixed at 42) and the initialisation yielding the highest log-likelihood was retained. Model complexity was penalised using the Bayesian Information Criterion:

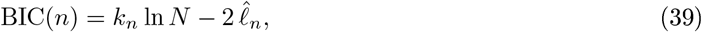

where *k*_*n*_ is the number of free parameters for the *n*-component model, *N* is the total number of observations, and 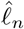 is the maximised log-likelihood. The model minimising Eq. (3) was selected as the best fit; a best-fit component count of *n*^∗^ = 2 was taken as evidence of bistability in the signal distribution.

#### 10.14.3 State dwell-time analysis

To characterise the temporal persistence of each oscillatory class (state), per-cell dwell times were extracted from the group-identity label sequence. Each cell was assigned a binary state label at every time point: state 0 for Class 1 cells and state 1 for Class 2 cells. Run-length encoding was applied to the label sequence of each cell to obtain a list of consecutive residence durations in each state. Dwell times were pooled across all cells and recordings within a condition for reporting.

The distribution of dwell times *τ* in each state was characterised by its mean, median, and maximum, and was compared against an exponential null model. An exponential distribution with rate parameter *λ* was fitted to the pooled dwell times by maximum likelihood estimation with location fixed at zero:

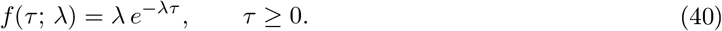

Exponentially distributed dwell times are consistent with thermally activated escape from a metastable potential well. Deviations from exponential behaviour — in particular heavy-tailed or multi-modal dwell-time distributions — would indicate more complex transition dynamics. Dwell-time histograms were plotted on a log-density scale to facilitate visual discrimination of exponential from non-exponential tails.

### 10.15 Low-Rank Recurrent Neural Network Model

To validate the geometric synchrony pipeline and provide a mechanistic interpretation of directed coupling between pituitary cell populations, a rank-2 recurrent neural network (RNN) was simulated under three coupling conditions: directed coupling from population A to B (*g*_*AB*_ *>* 0, *g*_*BA*_ = 0), null coupling (*g*_*AB*_ = *g*_*BA*_ = 0), and reversed coupling from B to A (*g*_*AB*_ = 0, *g*_*BA*_ *>* 0).

#### 10.15.1 Networ architecture and dynamics

Each population *P* ∈ {*A, B*} consists of *N*_*P*_ leaky-integrator units governed by:

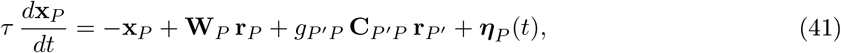

where 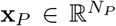 is the state vector, **r**_*P*_ = tanh(**x**_*P*_) is the firing-rate output, 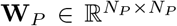 is the within-population recurrent weight matrix, 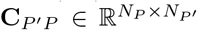 is the rank-1 cross-population coupling matrix, *g*_*P*_ *′*_*P*_ is the coupling gain, and ***η***_*P*_ (*t*) (**0**, ∼ *N σ* Δ*t* **I**) is additive Gaussian noise. Equation (41) was integrated with the Euler method using time step Δ*t* = 0.5 and membrane time constant *τ* = 5.0, giving an effective per-step update factor Δ*t/τ* = 0.1.

#### 10.15.2 Low-ran connectivity

The recurrent weight matrix of each population was constructed as a rank-*r* outer product:

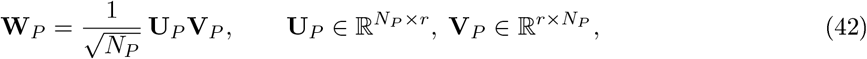

with *r* = 2. The matrix was then rescaled so that its dominant eigenvalue equals the target spectral radius *ρ* = 1.15, placing the network in the oscillatory (weakly chaotic) regime:

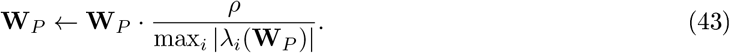

The cross-population coupling matrices were drawn as rank-1 random matrices, **C**_*AB*_ = ***ξζ***^⊤^*/N*_*A*_ with 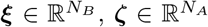 independently sampled from *N* (0, 1). Both populations used a shared random seed for **U, V**, and **C**, ensuring comparable geometric structure across conditions while the coupling gains *g*_*AB*_ and *g*_*BA*_ alone broke the symmetry.

#### 10.15.3 Simulation parameters

Populations contained *N*_*A*_ = 50 and *N*_*B*_ = 80 units. Each simulation ran for *T* = 1,200 time steps following random initialisation (**x**_*P*_ (0) ∼ *N* (**0**, 0.09 **I**)). Coupling gains were set to *g* = 3.0 for the active direction and *g* = 0 for the suppressed direction. Noise amplitude was *σ* = 0.015. All random seeds were fixed for reproducibility (weight seed = 8, initial condition seed = 108).

### 10.16 Geometric Synchrony Pipeline Applied to the RNN

The geometric synchrony pipeline described in precedent Section, was applied to the RNN trajectories with the following adaptations.

#### 10.16.1 Strict separation of lag estimation and geometric alignment

A key methodological constraint is that Procrustes alignment and cross-correlation lag estimation must not be chained. Applying Procrustes before cross-correlation erases the temporal lag by rotating PopB to maximally overlap PopA simultaneously across all time points, artificially inflating apparent synchrony. Therefore:

1. **Temporal lag** was estimated on sign-aligned raw PCA trajectories (**Z**_*A*_, sign(**Z**_*B*_)) via cross-correlation of the first three principal components, searching over both PCA sign polarities to resolve the sign ambiguity inherent to PCA. The peak lag across components and polarities was retained.
2. **Geometric metrics** (pointwise distance, subspace angles, sliding-window alignment) were computed after Procrustes alignment of **Z**_*B*_ onto **Z**_*A*_, as described in Section .

The lag convention adopted is: a negative lag (cross-correlation peak at *τ <* 0) indicates that *A* leads *B* (B is a delayed copy of A), while a positive lag indicates that *B* leads *A*.

#### 10.16.2 Validation criteria

A condition was considered correctly recovered if the peak lag satisfied: directed (*g*_*AB*_ = 3): *τ <* −3 (A leads); null: |*τ* | ≤ 15 (no systematic lead); reversed (*g*_*BA*_ = 3): *τ >* 3 (B leads).

### 10.17 Quasi-Potential Energy Landscape

To characterise the geometry of the population attractor and its evolution over time, a quasi-potential energy landscape was estimated from the latent trajectories. The landscape *U* was defined as the negative log-probability density of the population state in the first two latent dimensions:

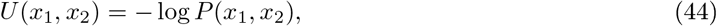

where *P* was estimated by a Gaussian kernel density estimator (KDE) with bandwidth parameter *h* = 0.35 (Scott’s method):

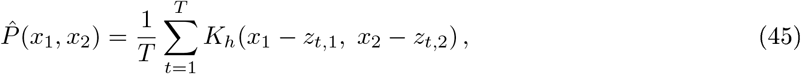

evaluated on an 80 × 80 grid spanning the observed latent range with 0.5-unit margins on each side. The estimated landscape was smoothed with a Gaussian filter (*σ* = 2.0 grid units) and shifted so that its minimum equals zero. Minima of *U* correspond to preferred (high-probability) population states; maxima correspond to energy barriers separating distinct attractor basins.

#### 10.17.1 Temporal landscape analysis

To capture the time-varying geometry of the shared attractor, the landscape was re-estimated within sliding windows of half-width *w*_*ℓ*_ = 30 samples, pooling the trajectories of both populations within each window. Two rolling metrics were derived:

1. **Well depth**: *U*_max_ − *U*_min_ within the window, quantifying attractor strength; deeper wells correspond to more confined, structured population dynamics.
2. **Centroid separation**: Euclidean distance between the time-windowed mean positions of PopA and PopB in latent space, serving as a proxy for geometric dominance (larger separation indicates that the two populations occupy distinct regions of the attractor).

Snapshot landscapes were also constructed for four equal temporal windows (each of duration *T/*4) and visualised as both three-dimensional surfaces (*U* versus latent dimensions 1 and 2) and two-dimensional contour maps with time-gradient coloured trajectories.

## References

1. Mollard, P., Hodson, D. J., Lafont, C., Rizzoti, K. & Drouin, J . A tridimensional view of pituitary development and function. Trends in Endocrinology & Metabolism 23, 261–269. ISSN: 1043-2760. https://www.sciencedirect.com/science/article/pii/S1043276012000288 (2025) (June 2012).

2. Le Tissier, P. et al. An updated view of hypothalamic-vascular-pituitary unit function and plasticity. en. Nature Reviews Endocrinology 13, 257–267. ISSN: 1759-5037. https://www.nature.com/articles/nrendo.2016.193 (2025) (May 2017).

3. Bonnefont, X. & Mollard, P. Electrical activity in endocrine pituitary cells in situ: A support for a multiple-function coding. FEBS Letters 548, 49–52. ISSN: 0014-5793. https://www.sciencedirect.com/science/article/pii/S0014579303007270 (2025) (July 2003).

4. Bonnefont, X., Fiekers, J., Creff, A. & Mollard, P. Rhythmic bursts of calcium transients in acute anterior pituitary slices. Endocrinology 141, 868–875. ISSN: 0013-7227 (Mar. 2000).

5. Hodson, D. J. et al. Existence of long-lasting experience-dependent plasticity in endocrine cell networks. en. Nature Communications 3, 605. ISSN: 2041-1723. https://www.nature.com/articles/ncomms1612 (2025) ( Jan. 2012).

6. Drummond, J. B., Ribeiro-Oliveira, A. & Soares, B. S. eng. in Endotext (eds Feingold, K. R.et al.) (MDText.com, Inc., South Dartmouth (MA), 2000). http://www.ncbi.nlm.nih.gov/books/NBK534880/ (2026).

7. Molitch, M. E. Pituitary incidentalomas. Best Practice & Research Clinical Endocrinology & Metabolism. Pituitary Tumours 23, 667–675. ISSN: 1521-690X. https://www.sciencedirect.com/science/article/pii/S1521690X09000487 (2026) (Oct. 2009).

8. Sanchez-Cardenas, C. et al. Pituitary growth hormone network responses are sexually dimorphic and regulated by gonadal steroids in adulthood. eng. Proceedings of the National Academy of Sciences of the United States of America 107, 21878–21883. ISSN: 1091-6490 (Dec. 2010).

9. Bonnefont, X. et al. Revealing the large-scale network organization of growth hormone-secreting cells. Proceedings of the National Academy of Sciences 102, 16880–16885. 10.1073/pnas.0508202102 (2025) (Nov. 2005).

10. Fauquier, T., Guérineau, N. C., McKinney, R. A., Bauer, K. & Mollard, P. Folliculostellate cell network: A route for long-distance communication in the anterior pituitary. Proceedings of the National Academy of Sciences 98, 8891–8896. https://www.pnas.org/doi/abs/10.1073/pnas.151339598 (2025) (July 2001).

11. Le Tissier, P. R. et al. Anterior pituitary cell networks. eng. Frontiers in Neuroendocrinology 33, 252–266. ISSN: 1095-6808 (Aug. 2012).

12. Veldhuis, J. D., Carlson, M. L. & Johnson, M. L. The pituitary gland secretes in bursts: appraising the nature of glandular secretory impulses by simultaneous multiple-parameter deconvolution of plasma hormone concentrations. Proceedings of the National Academy of Sciences of the United States of America 84, 7686–7690. ISSN: 0027-8424. https://pmc.ncbi.nlm.nih.gov/articles/PMC299365/ (2026) (Nov. 1987).

13. Goor, F. V., Zivadinovic, D., Martinez-Fuentes, A. J. & Stojilkovic, S. S. Dependence of Pituitary Hormone Secretion on the Pattern of Spontaneus Voltage-gated Calcium Influx. English. Journal of Biological Chemistry 276, 33840–33846. ISSN: 0021-258, 1083-351. https://www.jbc.org/article/S0021-9258(19)34785-4/abstract (2025) (Sept. 2001).

14. Stojilkovic, S. S. Pituitary cell type-specific electrical activity, calcium signaling and secretion. Biological Research 39, 403–423. ISSN: 0716-760. http://www.scielo.cl/scielo.php?script=sci_abstract&pid=S0716-97602006000300004&lng=es&nrm=iso&tlng=en (2025) (2006).

15. Stojilkovic, S. S. Signaling pathways regulating pituitary functions. Molecular and cellular endocrinology 463, 1–3. ISSN: 0303-7207. https://www.ncbi.nlm.nih.gov/pmc/articles/PMC6326166/ (2025) (Mar. 2018).

16. Christian, H. C. en. in Neurosecretion: Secretory Mechanisms (eds Lemos, J. R. & Dayanithi, G.) 173–1 3 (Springer International Publishing, Cham, 2020). ISBN: 978-3-030-22989-4. 10.1007/978-3-030-22989-4_9 (2026).

17. Santiago-Andres, Y., Golan, M. & Fiordelisio, T. Functional Pituitary Networks in Vertebrates. Frontiers in Endocrinology 11. ISSN: 1664-2392. https://www.frontiersin.org/articles/10.3389/fendo.2020.619352 (2023) (2021).

18. Briano, C., Meikle, A., Velazco, J. I. & Quintans, G. Metabolic and hormonal profiles and productive performance in primiparous and multiparous cows grazing different forage allowance in late gestation. Theriogenology 227, 68–76. ISSN: 0093-691X. https://www.sciencedirect.com/science/article/pii/S0093691X24002735 (2026) (Oct. 2024).

19. en-US. in Maternal-Fetal and Neonatal Endocrinology 189-205 (Academic Press, Jan. 2020). https://www.sciencedirect.com/science/chapter/edited-volume/abs/pii/B9780128148235000143 (2026).

20. Brake, N. et al. A neurophysiological basis for aperiodic EEG and the background spectral trend. en. Nature Communications 15, 1514. ISSN: 2041-1723. https://www.nature.com/articles/s41467-024-45922-8 (2024) (Feb. 2024).

21. Donoghue, T. A historical overview of the study of aperiodic neural activity. Nov. 2024. https://osf.io/zrvxa_v1 (2025).

22. Donoghue, T. A systematic review of aperiodic neural activity in clinical investigations. Pages: 2024.10.14.24314 25. Oct. 2024. https://www.medrxiv.org/content/10.1101/2024.10.14.24314925v1 (2025).

23. Aquiles, A., Pinedo-Vargas, A. L., Luna-Munguía, H. & Rossi-Pool, R. Assessing the degree of cortical dislamination through electrical pattern analysis. iScience 29, 115942 (2026).

24. Rombouts, J. & Gelens, L. Dynamic bistable switches enhance robustness and accuracy of cell cycle transitions. P LoS Computational Biology 17, e1008231. ISSN: 1553-734X. https://pmc.ncbi.nlm.nih.gov/articles/PMC7817062/ (2026) (Jan. 2021).

25. Joshi, B. & Shiu, A. A Survey of Methods for Deciding Whether a Reaction Network is Multistationary. en. Mathematical Modelling of Natural Phenomena 10, 47–67. ISSN: 0973-5348, 1760-6101. https://www-mmnp-journal-org.insb.bib.cnrs.fr/articles/mmnp/abs/2015/05/mmnp201510p47/mmnp201510p47.html (2026) (2015).

26. Strogatz, S. H. & Stewart, I. Coupled Oscillators and Biological Synchronization. Scientific American 269, 102–109 . ISSN: 0036-8733. https://www.jstor.org/stable/24941731 (2023) (1993).

27. Izhikevich, E. M. Dynamical Systems in Neuroscience: The Geometry of Excitability and Bursting en. ISBN: 978-0-262-27607-8. https://direct.mit.edu/books/monograph/2589/Dynamical-Systems-in-NeuroscienceThe-Geometry-of (2026) (The MIT Press, July 2006).

28. Mastrogiuseppe, F. & Ostojic, S. Linking Connectivity, Dynamics, and Computations in Low-Rank Recurrent Neural Networks. Neuron 99, 609-623.e29 . ISSN: 0896-6273. https://www.sciencedirect.com/science/article/pii/S0896627318305439 (2024) (Aug. 2018).

29. Izhikevich, E. M. Dynamical Systems in Neuroscience. ISBN: 978-0-262-09043-8 (MIT Press, 2007).

30. Le Tissier, P. et al. An updated view of hypothalamic-vascular-pituitary unit function and plasticity. eng. Nature Reviews. Endocrinology 13, 257–267. ISSN: 1759-5037 (May 2017).

31. Spontaneous oscillations of intracellular calcium and growth hormone secretion. en-US. Journal v 263, 9682–9685. ISSN: 0021-9258. https://www.sciencedirect.com/science/article/pii/S0021925819815715 (2025) (July 1988).

32. Kidokoro, Y. Spontaneous calcium action potentials in a clonal pituitary cell line and their relationship to prolactin secretion. en. Nature 258, 741–742. ISSN: 1476-4687. https://www.nature.com/articles/258741a0 (2025) (Dec. 1975).

33. Kwiecien, R. & Hammond, C. Differential Management of Ca2+ Oscillations by Anterior Pituitary Cells: A Comparative Overview. Neuroendocrinology 68, 135–151. ISSN: 0028-3835. 10.1159/000054360 (2026) (Sept. 1998).

34. Kuryshev, Y. A., Childs, G. V. & Ritchie, A. K. Corticotropin-releasing hormone stimulates Ca2+ entry through L- and P-type Ca2+ channels in rat corticotropes. Endocrinology 137, 2269-2277. ISSN: 0013-7227. https://doi.org/10.1210/en.137.6.2269 (2026) (June 1996).

35. Tsaneva-Atanasova, K., Sherman, A., van Goor, F. & Stojilkovic, S. S. Mechanism of spontaneous and receptor-controlled electrical activity in pituitary somatotrophs: experiments and theory. eng. V 98, 131–144. ISSN: 0022-3077 (July 2007).

36. Tabak, J., Tomaiuolo, M., Gonzalez-Iglesias, A. E., Milescu, L. S. & Bertram, R. Fast-activating voltage- and calcium-dependent potassium (BK) conductance promotes bursting in pituitary cells: a dynamic clamp study. eng. The Journal of Neuroscience: The Official Journal of the Society for Neuroscience 31, 16855–16863. ISSN: 1529-2401 (Nov. 2011).

37. Stojilkovic, S. S., Tabak, J. & Bertram, R. Ion Channels and Signaling in the Pituitary Gland. Endocrine Reviews 31, 845–915. ISSN: 0163-769X. https://doi.org/10.1210/er.2010-0005 (2025) (Dec. 2010).

38. Gahete, M. D. et al. Understanding the multifactorial control of growth hormone release by somatotropes: lessons from comparative endocrinology. eng. Annals of the New York Academy of Sciences 1163, 137–153. ISSN: 1749-6632 (Apr. 2009).

39. Gao, R., Peterson, E.. & Voytek, B. Inferring synaptic excitation/inhibition balance from field potentials. NeuroImage 158, 70–78. ISSN: 1053-8119 . https://www.sciencedirect.com/science/article/pii/S1053811917305621 (2025) (Sept. 2017).

40. Donoghue, T. et al. Parameterizing neural power spectra into periodic and aperiodic components. en. Nature Neuroscience 23, 1655–1665. ISSN: 1546-1726. https://www.nature.com/articles/s41593-020-00744-x (2025) (Dec. 2020).

41. Voytek, B. et al. Age-Related Changes in 1/f Neural Electrophysiological Noise. eng. The Journal of Neuroscience: The Official Journal of the Society for Neuroscience 35, 13257–13265. ISSN: 1529-2401 (Sept. 2015).

42. He, M.-L., Gonzalez-Iglesias, A. & Stojilkovic, S. Role of Nucleotide P2 Receptors in Calcium Signaling and Prolactin Release in Pituitary Lactotrophs. The Journal of biological chemistry 278, 46270–7 (Dec. 2003).

43. Stojilkovic, S. S., Bjelobaba, I. & Zemkova, H. Ion Channels of Pituitary Gonadotrophs and Their Roles in Signaling and Secretion. Frontiers in Endocrinology 8, 126. ISSN: 1664-2392. https://pmc.ncbi.nlm.nih.gov/articles/PMC5465261/ (2026) (June 2017).

44. Sneyd, J. et al. On the dynamical structure of calcium oscillations. Proceedings of the National Academy of Sciences 114. Publisher: Proceedings of the National Academy of Sciences, 1456–1461. https://www.pnas.org/doi/10.1073/pnas.1614613114 (2026) (Feb. 2017).

45. Bertram, R., Tabak, J., Teka, W., Vo, T. & Wechselberger, M. en. in Mathematical Analysis of Complex Cellular Activity (eds Bertram, R.et al.) 1–52 (Springer International Publishing, Cham, 2015). ISBN: 978-3-31 9-18114-1. https://doi.org/10.1007/978-3-319-18114-1_1 (2026).

46. Fletcher, P. A. & Li, Y.-X. An Integrated Model of Electrical Spiking, Bursting, and Calcium Oscillations in GnRH Neurons. English. Biophysical Journal 96. Publisher: Elsevier, 4514–4524. ISSN: 0006-3495, 1542-0086. https://www.cell.com/biophysj/abstract/S0006-3495(09)00757-7 (2026) (mJune 2009).

47. Gallego, J. A., Perich, M. G., Miller, L. E. & Solla, S. A. Neural Manifolds for the Control of Movement. eng. Neuron 94, 978–984. ISSN: 1097-4199 (June 2017).

48. Rossi-Pool, R. & Romo, R. Low Dimensionality, High Robustness in Neural Population Dynamics. English. Neuron 103. Publisher: Elsevier, 177–179 . ISSN: 0896-6273. https://www.cell.com/neuron/abstract/S0896-6273(19)30571-9 (2026) (July 2019).

49. Virtanen, P. et al. SciPy 1.0: fundamental algorithms for scientific computing in Python. en. Nature Methods 17. Number: 3, 261–272. ISSN: 1548-7105. https://www.nature.com/articles/s41592%E2%80%90019%E2%80%900686%E2%80%902 (2023) (Mar. 2020).

50. Tort, A. B. L., Komorowski, R., Eichenbaum, H. & Kopell, N. Measuring phase-amplitude coupling between neuronal oscillations of different frequencies. eng. Journal of Neurophysiology 104, 1195–1210. ISSN: 1522-1598 (Aug. 2010).

51. Pedregosa, F. et al. Scikit-learn: Machine learning in Python. the Journal of machine Learning research 12, 2825–2830. http://www.jmlr.org/papers/volume12/pedregosa11a/pedregosa11a.pdf?source=post_page (2025) (2011).

52. Aquiles, A. et al. Assessing the degree of cortical dislamination through electrical pattern analysis en. Pages: 2025.06.16.659942 Section: New Results. June 2025. https://www.biorxiv.org/content/10.1101/2025.06.16.659942v2 (2025).

53. Neville, M. C., McFadden, T. B. & Forsyth, I. Hormonal regulation of mammary differentiation and milk secretion. Journal of Mammary Gland Biology and Neoplasia 7, 49–66 (Jan. 2002).

54. Mena, F., Enjalbert, A., Carbonell, L., Priam, M. & Kordon, C. Effect of suckling on plasma prolactin and hypothalamic monoamine levels in the rat. Endocrinology 99, 445–451 (Aug. 1976).

55. Grattan, D. R., Steyn, F. J., Kokay, I. C., Anderson, G. M. & Bunn, S. J. Pregnancy-Induced Adaptation in the Neuroendocrine Control of Prolactin Secretion. Journal of Neuroendocrinology 20, 497–507 (2008).

56. Anderson, G. M., Grattan, D. R., van den Ancker, W. & Bridges, R. S. Reproductive experience increases prolactin responsiveness in the medial preoptic area and arcuate nucleus of female rats. Endocrinology 147, 4688–4694 (Oct. 2006).

57. Haggi, E. S., Torres, A. I., Maldonado, C. A. & Aoki, A. Regression of redundant lactotrophs in rat pituitary gland after cessation of lactation. Journal of Endocrinology 111, 367–373 (Dec. 1986).

